# A self-amplifying mRNA COVID-19 vaccine drives potent and broad immune responses at low doses that protects non-human primates against SARS-CoV-2

**DOI:** 10.1101/2021.11.08.467773

**Authors:** Amy R. Rappaport, Sue-Jean Hong, Ciaran D. Scallan, Leonid Gitlin, Arvin Akoopie, Gregory R. Boucher, Milana Egorova, J. Aaron Espinosa, Mario Fidanza, Melissa A. Kachura, Annie Shen, Gloria Sivko, Anne Van Abbema, Robert L. Veres, Karin Jooss

## Abstract

The coronavirus disease 2019 (COVID-19) pandemic continues to spread globally, highlighting the urgent need for safe and effective vaccines that could be rapidly mobilized to immunize large populations. We report the preclinical development of a self-amplifying mRNA (SAM) vaccine encoding a prefusion stabilized severe acute respiratory syndrome coronavirus 2 (SARS-CoV-2) spike glycoprotein and demonstrate potent cellular and humoral immune responses at low doses in mice and rhesus macaques. The homologous prime-boost vaccination regimen of SAM at 3, 10 and 30 μg induced potent neutralizing antibody titers in rhesus macaques following two SAM vaccinations at all dose levels, with the 10 μg dose generating geometric mean titers (GMT) 48-fold greater than the GMT of a panel of SARS-CoV-2 convalescent human sera. Spike-specific T cell responses were observed at all dose levels. SAM vaccination provided protective efficacy against SARS-CoV-2 challenge as both a homologous prime-boost and as a single boost following ChAd prime, demonstrating reduction of viral replication in both the upper and lower airways. Protection was most effective with a SAM prime-boost vaccination regimen at 10 and 30 μg and with a ChAd/SAM heterologous prime-boost regimen. The SAM vaccine is currently being evaluated in clinical trials as both a homologous prime-boost regimen at low doses and as a boost following heterologous prime.

## Introduction

The current severe acute respiratory syndrome coronavirus 2 (SARS-CoV-2) pandemic has spurred the rapid development and approval of multiple vaccines. Yet, despite this unprecedented rate of response, the virus continues to spread globally with a devasting humanitarian toll (> 4.8 million global deaths, approx. 236 million infections as of October 6^th^, 2021) (https://coronavirus.jhu.edu/map.html) and major economic impact. The pandemic highlights disparities in vaccine availability, with the developing world being disproportionally impacted; fewer than 1% of people in low-income countries are fully vaccinated against SARS-CoV-2 compared to >50% in high income countries^1^. Delays in global vaccination allow the virus to continue to spread and mutate, leading to the emergence of new variants of concern (VOC), such as the Delta variant (B.1.617.2). Additional vaccine doses are needed which could be achieved not only by expanding manufacturing capacity but also by advancing new vaccine platforms, such as self-amplifying mRNA (SAM) vaccines, which could potentially be dose sparing due to the platform’s ability to replicate post vaccination.

Two main vaccine platforms have come to the fore in this pandemic, mRNA (mRNA-1273, Moderna; BNT162b2, Pfizer/BioNTech) and adenovirus based vaccines (ChAdOx1/AZD1222, AstraZeneca; Ad26.COV2.S, Janssen), both of which have demonstrated potent protection from hospitalization and death^2-5^. Adenovirus (Ad) based vaccine vectors have a long history of development and have been shown to be potent inducers of both humoral and cellular immunity, with an Ad26 based vaccine approved for prevention of Ebola by the European Commission (https://ec.europa.eu/commission/presscorner/detail/en/ip_20_1248). The authorized adenovirus COVID vaccines demonstrate vaccine effectiveness in the range of 64.3% to 71%^4,6^. Vascular induced thrombotic thrombocytopenia (VITT), a rare clotting issue, has been associated with the Ad-SARS-CoV-2 vaccines^7^, resulting in some countries preferentially offering mRNA vaccines, especially to younger and female populations.

mRNA based vaccines are relatively new to the clinic, with the Pfizer/BioNTech BNT162b2 and Moderna mRNA-1273 COVID vaccines being the first to receive emergency use authorization. Both have demonstrated potency in pre-clinical and clinical testing^2,3,8,9^, with a standard vaccination regimen of two doses given 3-4 weeks apart to drive optimal immune responses. The authorized mRNA COVID vaccine doses are either 30 μg for the BNT162b2 vaccine (Pfizer/BioNTech) or 100 μg for mRNA-1273 (Moderna), and both provide clinical protection in the range of 88-93%^6^. Production of mRNA vaccines by a cell-free enzymatic transcription reaction *in vitro* enables simple downstream purification and rapid vaccine release, providing for a cost-effective manufacturing process. Non-viral delivery systems, such as lipid nanoparticles (LNPs), allow repeated RNA administrations without inducing neutralizing anti-vector immunity.

SAM vaccines offer the potential benefit of driving equivalent or more potent immune responses at lower doses compared to non-replicating mRNA vaccines, as the mRNA replicates within the cell, leading to the activation of innate signaling pathways and strong and durable antigen expression^10^. We have developed and tested SAM based vaccines, initially in immuno-oncology applications, and have demonstrated their safety and potency in boosting neoantigen-specific T-cell responses primed by a chimpanzee adenovirus (ChAd68) vaccine in human clinical trials (manuscript in preparation).

In this study, we evaluated a SAM-SARS-CoV-2 vaccine either as a homologous prime/boost vaccine regimen (SAM/SAM), or as a boost vaccine following a ChAd-SARS-CoV-2 prime (ChAd/SAM) in both mice and non-human primates (NHP). The SAM/SAM homologous prime/boost regimen allows comparison with the authorized mRNA COVID-19 vaccines, while the ChAd/SAM vaccine regimen was assessed to study the potency of a COVID-19 vaccine regimen combining ChAd and mRNA-based vaccines in a heterologous prime/boost approach. The latter mix-and-match approach has been evaluated clinically and offered to individuals who had initially received the ChAdOx1 (AZD1222/AstraZeneca) or Ad26.COV2.S (Janssen) vaccines^11,12^.

While developing the ChAd and SAM COVID vaccine platforms, we observed that the ChAd vaccine required extensive spike sequence optimization to drive potent neutralizing antibody responses, a step that is beneficial but not necessary for SAM. In rhesus macaques, a SAM/SAM homologous prime-boost regimen elicited robust neutralizing antibody titers that were comparable to immune responses generated by the ChAd/SAM heterologous prime-boost regimen. While ChAd prime generated increased and more rapid T cell responses as a single dose, T cell response following SAM boost were similar with both heterologous and homologous regimens. Furthermore, the protective immunity induced by SAM/SAM vaccination, which appeared to be equivalent to currently authorized conventional mRNA vaccines, was achieved at lower doses.

Taken together, the data demonstrate that the SAM vaccine platform can drive balanced T- and B-cell responses for optimal protective immunity at low doses, offering an additional attractive vaccine platform in the fight against the ongoing SARS-CoV-2 pandemic and against current and emerging infectious pathogens. The SAM SARS-CoV-2 vaccine is currently being evaluated in multiple clinical trials as both a homologous prime/boost regimen or as a boost in adenovirus primed individuals.

## Results

### Codon optimization improves spike expression and immunogenicity

SAM and ChAd vaccine vectors encoding a full-length codon optimized SARS-CoV-2 spike sequence (V1) were evaluated for immunogenicity in mice. While the ChAd-Spike(V1) vaccine vector induced detectable neutralizing antibody titers in 4/8 mice 8-weeks post immunization (pseudovirus 50% inhibitory titer (NT50) geomean titer (GMT) 40) (Figure 1B) and a strong spike-specific T cell response (Mean ± SE 5,246 ± 363 SFU/10^6^ splenocytes) 2-weeks post immunization (Figure 1C), similar to those observed with other adenoviral spike vectors^13^, the spike-specific immune response induced by the SAM-Spike(V1) vaccine was significantly higher in all (12/12) vaccinated animals (NT50 GMT 396 & IFNγ T cell response 9,793 ± 1,067 SFU/10^6^ splenocytes) than the response observed with the ChAd vaccine (Figure 1B&C). We had not previously observed significantly stronger immune responses induced by the SAM vector compared to the ChAd vector in mice using multiple antigens (data not shown), prompting a spike sequence optimization campaign to increase the ChAd-Spike vaccine potency. Initial data with the ChAd-Spike construct indicated a low level of spike protein expression by Western blot in vitro (data not shown). We hypothesized that nuclear processing, transport, or stability, as required by DNA and not RNA-based vectors, impacted overall expression levels. Therefore, optimization efforts focused on improving spike expression driven by the ChAd vectored Spike(V1) protein. Seven additional codon optimized sequences were generated using two different codon optimization tools. The resulting ChAd-Spike viral vectors were assessed for spike expression by Western blot analyses specific to spike S2. A band of 170 kDa, corresponding to full-length spike protein, was the major form of the expressed protein. Expression was highly variable across different codon optimized spike sequences, with V2 and V8 having strong expression, V4 and V6 having intermediate levels of expression and V1, V3, V5 and V7 demonstrating weak expression (Figure 1A). Consistent with increased expression *in vitro*, the ChAd-Spike(V2) vaccine induced significantly higher neutralizing antibody titers in mice compared with Spike(V1) (NT50 GMT 2,525 vs. 40, 63-fold) with a trend toward increased T cell response (1.7-fold). In contrast, a SAM-Spike(V2) vaccine resulted in lesser improvements in B and T cell responses compared with SAM-Spike(V1), but similar immune response to that induced by the ChAd-Spike(V2) vaccine (Figure 1B&C). Both ChAd & SAM Spike(V2) vaccines demonstrated increased durability of total spike IgG antibody titers as compared with Spike(V1) (Supp. Fig. 1). Potent immune response was observed *in vivo* with the Spike(V8) variant (data not shown), which also correlated with robust expression *in vitro* (Figure 1A). These results demonstrate that sequence optimization can significantly impact antigen expression and vaccine potency differentially across vaccine platforms.

**Figure 1.**
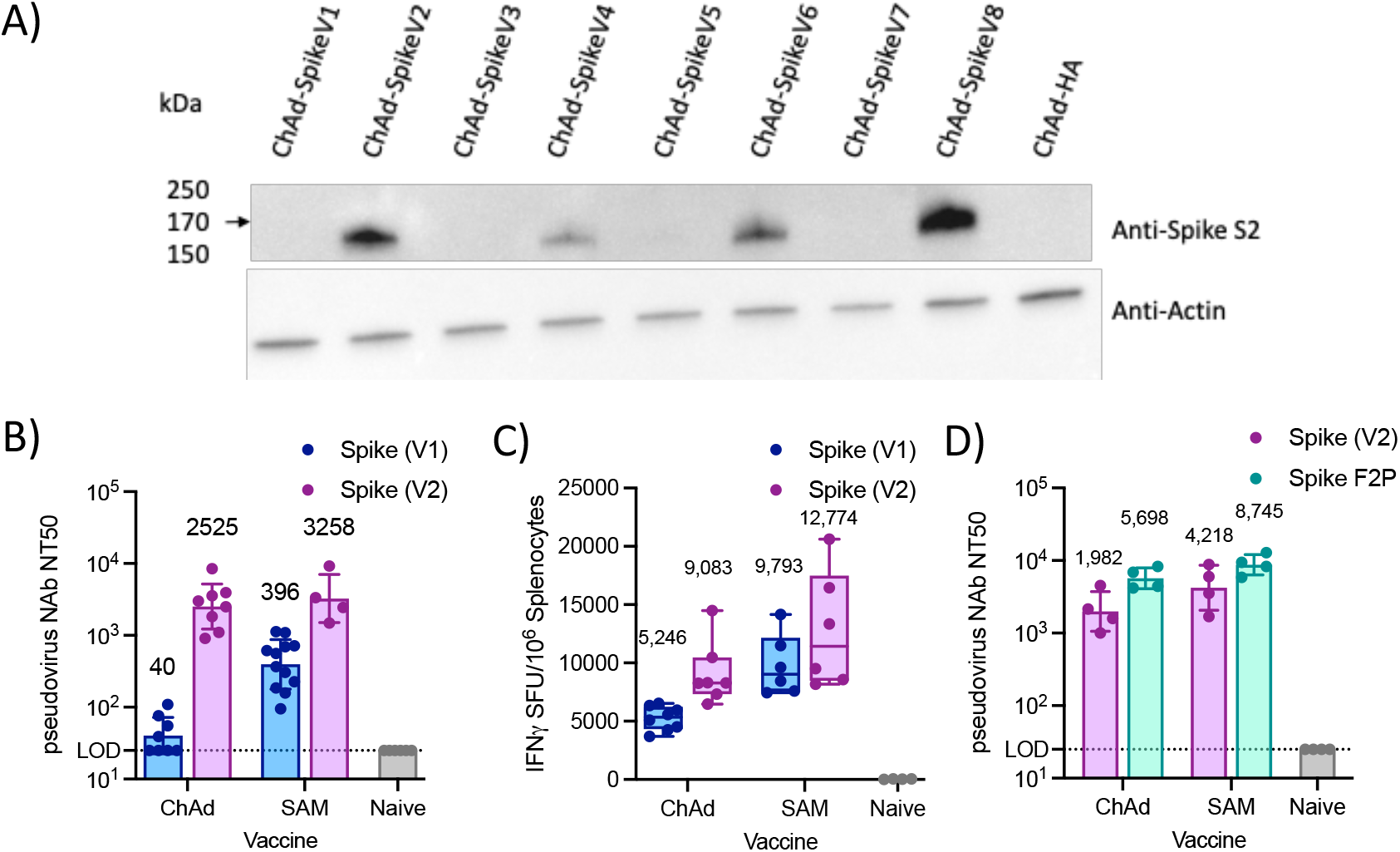
Codon optimization of spike sequence increases antigen expression *in vitro* and immune response in mice. (A) Expression analysis of eight codon optimized spike sequences (V1-V8) expressed using ChAd vaccine vectors, assessed by Western blot 40 hours post infection in HEK293 cells using an anti-spike S2 antibody. A ChAd-influenza HA construct served as a negative control. An anti-Actin antibody was used as loading control. (B) Serum pseudovirus neutralizing antibody titers (50% inhibition) in Balb/c mice 8 weeks post immunization with either ChAd (1×10^11^ VP) or SAM (10 μg) encoding either the spike V1 or V2 sequence. Naïve samples all below LOD (LOD = 25). Geomean and geometric SD. Numbers are GMT. (C) Antigen-specific T-cell response assessed by IFNγ ELISpot (sum of two spike pools) in splenocytes in Balb/c mice 2 weeks post immunization with either ChAd (1×10^11^ VP) or SAM (10 μg) encoding either the Spike V1 or V2 sequence. Box and whiskers represent median, IQR and range. Numbers are mean. Note that ChAd and SAM vaccines were assessed in separate studies. (D) Serum pseudovirus neutralizing antibody titers (50% inhibition) in Balb/c mice 8 weeks post immunization with ChAd (1×10^11^ VP) or SAM (10 μg) encoding the specified spike variant. Geomean and geometric SD. Numbers are GMT. Naïve mice all below LOD (LOD = 25).

### Prefusion stabilized spike improves humoral immunity

To further improve vaccine potency, vectors expressing the prefusion-stabilized spike protein^14^ were generated. The Spike(V2) sequence was modified by mutation of the furin cleavage (FurinΔ) site between amino acids 681 and 685 and introduction of proline substitutions to the spike S2 domain (2P: K986P, V987P)^15^ (F2P) (Supp. Fig. 2A). This F2P variant demonstrated increased neutralizing antibody titers compared with V2 for both ChAd (NT50 GMT 5,698 vs 1,982, 2.9-fold) and SAM (NT50 GMT 8,745 vs 4,218, 2.1-fold), with more substantial benefit provided to the ChAd vaccine as compared to the SAM vaccine (Figure 1D, Supp. Fig. 2G). Additional vectors containing alternate furin modifications and proline substitutions also demonstrated increased antibody titers compared with V2 (Supp. Fig. 2D,E). T cell response remained potent but unchanged by the spike modifications (Supp. Fig. 2C,F).

**Figure 2.**
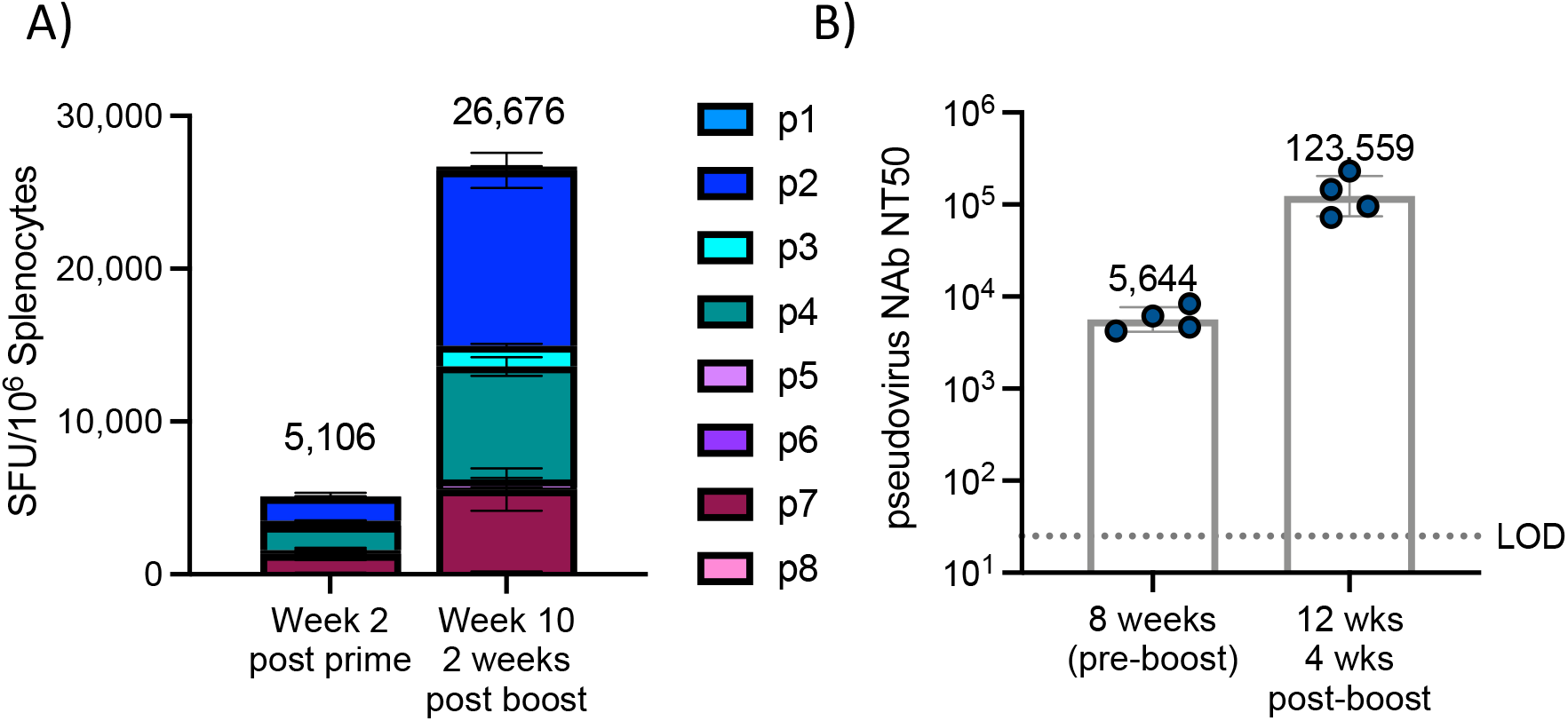
SAM boost immunization drives increased antigen specific T cells and neutralizing antibody titers. Mice received SAM prime immunization, followed by boost 8 weeks post prime (10 μg each). N = 4/group. (A) IFNγ ELISpot at the specified timepoint, following stimulation with eight peptide pools spanning spike antigen. Mean ± SEM for each peptide pool. (B) Pseudovirus neutralizing titer (NT50) at the specified timepoint. Geometric mean and geometric SD. Sera from Naïve mice were below LOD of 25.

### SAM homologous prime/boost regimen drives potent immune response in mice

Optimized Spike(V2)-F2P ChAd and SAM vaccines were evaluated for immunogenicity in Balb/c mice, as either a homologous prime/boost (SAM prime and SAM boost) or a heterologous prime/boost (ChAd prime and SAM boost), with an eight-week interval between prime and boost. Both vaccines generated potent and broad spike-specific T cell responses, which were increased following SAM boost immunization (2.5 and 5.2-fold) and were predominantly CD8^+^ and T_H_1-biased (Figure 2A; Supp. Fig. 3&4). Following prime immunization with either ChAd or SAM, potent neutralizing antibody titers were observed, which were increased following boost immunization with 10 μg of SAM (36-fold for heterologous and 22-fold for homologous) (Figure 2B, Supp. Fig. 3B).

### Self-amplifying mRNA (SAM) vaccine drives potent immune response in rhesus macaques at low doses

The robust immune response induced by SAM in mice, even at low doses (Supp. Fig. 7), was explored further in NHP. Groups of 5 rhesus macaques were immunized intramuscularly twice with SAM-Spike(V2)-F2P (4-week interval) at either 3, 10 or 30 μg (Figure 3A). Spike S1-specific IgG antibody titers were observed in some NHP following a single SAM immunization in a dose dependent manner and were increased to high levels in all NHP following a 2^nd^ SAM immunization at all doses tested (GMT 8,880, 13,145 and 1,223 at 30, 10 and 3 μg, respectively, Figure 3B). Spike-specific T cell responses, assessed by overnight IFNγ ELISpot using a single spike peptide pool, were detected at all dose levels following two immunizations (Figure 3C). T cell response was significantly increased compared to unvaccinated control animals for all 3 dose groups (p < 0.05 for all 3 dose groups by Mann-Whitney comparison of peak T cell values) and were not significantly different between the dose groups. ELISpot analysis demonstrated strong IFNγ, but minimal IL-4, responses one-week following the boost immunization in all vaccinated animals, indicating a T_H_1-biased response (Figure 3D). Strong neutralizing antibody titers were detected following two immunizations of SAM by both a pseudovirus neutralization assay (PNA) and live virus microneutralization assay (MNA), which strictly correlated (Figure 3E,F; Supp. Fig. 5). High and similar neutralizing antibody titers were detected in all NHP (10/10) at both the 30 and 10 μg dose level (NT50 GMT 2,180 and 2,590 2-weeks post boost by MNA), demonstrating comparable immune response with lower doses of SAM. Lowering the dose of SAM further to 3 μg resulted in overall lower, but clearly detectable neutralizing antibody titers in 5/5 NHP (by MNA) compared with the 10 and 30 μg dose (GMT 245 2-weeks post boost) (Figure 3F). In addition, while potent spike-specific immune responses were observed at all dose levels, there was no increase in serum IFNα levels at 8 hours following each SAM immunization at doses ≤ 10 μg, suggesting that innate immune pathways are not activated at lower doses of SAM (Supp. Fig. 8)

**Figure 3.**
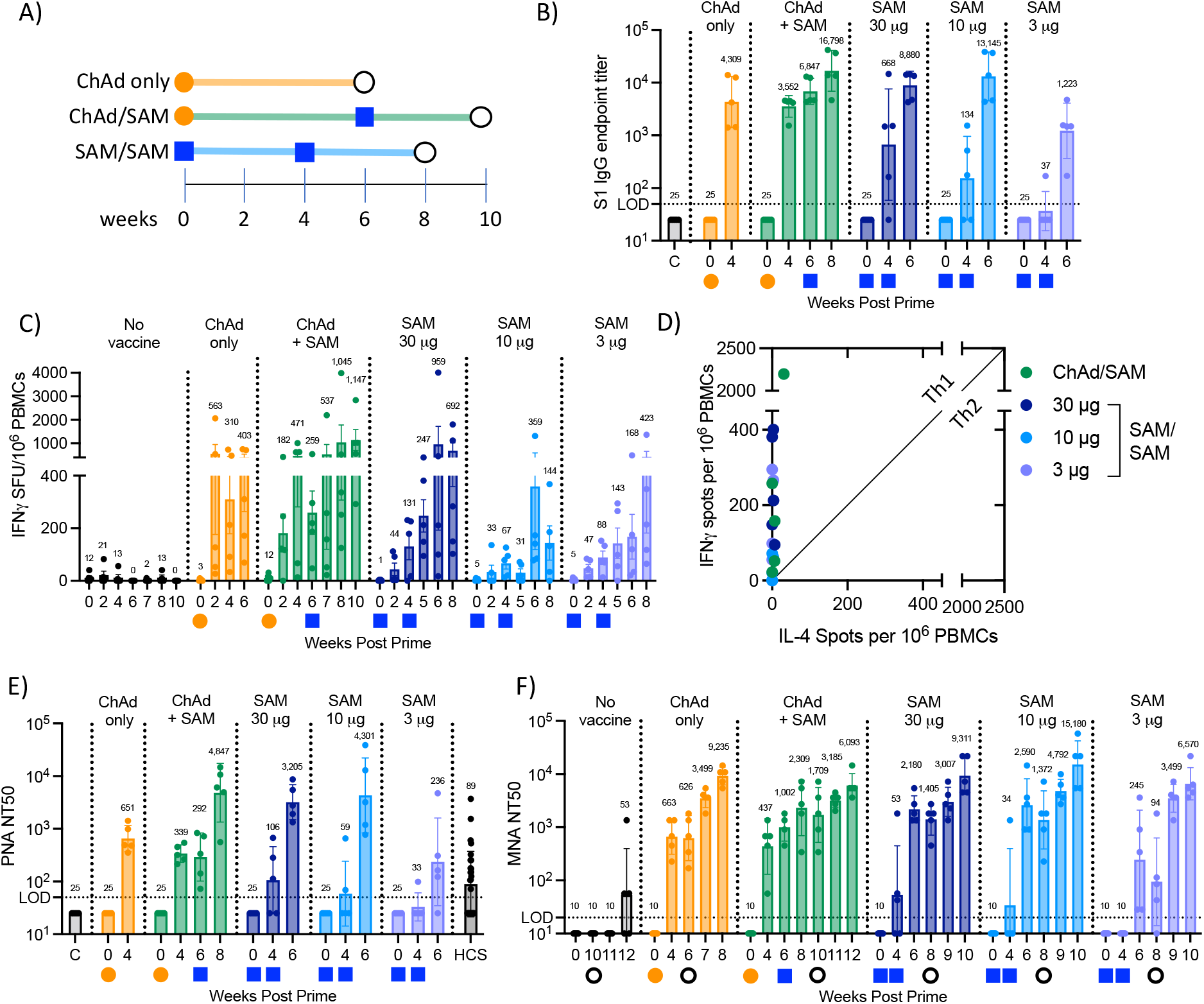
Immunogenicity in rhesus macaques. (A) Schematic of SARS-CoV-2 vaccine evaluation and challenge in rhesus macaques. Orange circles represent ChAd immunization (5×10^11^ VP), blue squares represent SAM vaccination at varying doses and black circle represents challenge with SARS-CoV-2 virus. N = 5/group. (B) spike S1 IgG endpoint titers, geometric mean (annotated) and geometric SD. LOD = 50, samples below LOD set to ½ LOD. (C) IFNγ ELISpot (SFU/10^6^ PBMCs) at specified timepoint post immunization, following overnight stimulation with overlapping peptide pool spanning spike antigen. Mean (annotated) +/-SEM. (D) IL-4 vs IFNγ ELISpot of PBMCs assessed 1-week post boost immunization. Individual animal values (n=5/group). Background corrected values. Diagonal line represents unity line. (E) Pseudovirus neutralization titers (50% inhibition, NT50) assessed in sera at specified timepoint post immunization. LOD = 50, samples < LOD set to ½ LOD. Geometric mean (annotated) and SD. HCS = human convalescent serum assessed by same assay. C = PBS control injected animals assessed 8 weeks post injection. (F) Live virus micro-neutralizing titer (50% inhibition, NT50) at specified timepoint post immunization and post SARS-CoV-2 challenge, geometric mean (annotated) and geometric SD. LOD = 20, samples below LOD set to ½ LOD.

### Low doses of SAM vaccine provide protection from SARS-CoV-2 infection

At 4-weeks following the 2^nd^ SAM immunization (week 8) (Figure 3A), all rhesus macaques were challenged with ∼1.6×10^6^ plaque-forming units of SARS-CoV-2 (strain USA-WA1/2020), split equally between the intratracheal and intranasal routes. Consistent with previous studies of SARS-CoV-2 challenge in rhesus macaques, none of the vaccinated or control animals showed clinical signs of illness^8,9,16-18^. Neutralizing antibody titers were detected in 3/5 phosphate-buffered saline (PBS) injected (control) animals 2-weeks post challenge (NT50 GMT 53). An increase in titers was observed 1- or 2-weeks post challenge in all SAM vaccinated animals, to much higher levels than those observed in control animals (NT50 GMT 9,311, 15,180 and 6,570 for SAM at 30, 10, and 3 μg, respectively, at 2-weeks post challenge), indicating an anamnestic response (Figure 3F).

To assess protective efficacy provided by the vaccine, viral load was assessed by PCR in bronchial alveolar lavage (BAL) fluid, oropharyngeal swabs, and nasal swabs at various times post challenge. Both total genomic RNA and subgenomic RNA (sgRNA) were assessed to quantify input virus and replicating virus, respectively. Three days after challenge, only one of five animals at the 30 μg and 3 μg dose levels and zero of five animals in the 10 μg group had detectable subgenomic RNA in BAL fluid, as compared to five of five animals in the control group. By day 7, none of the animals in any vaccine treated group had detectable sgRNA in the BAL, as compared to 2/5 in the control group, demonstrating protection from viral replication at all dose levels (Figure 4A). Comparison of peak viral titer for each animal between day 3 and day 10 post challenge by Mann-Whitney analysis demonstrated significant protection from viral replication at all dose levels compared to control animals (p = 0.008(**), 0.008(**) and 0.016(*) at 30, 10 and 3 μg, respectively, Supp. Fig. 9). In the oropharyngeal swab at day 2, none of the animals in the 10 μg dose group had detectable sgRNA, compared to one of five in the 30 μg SAM group, three of five in the control group and 3 μg SAM group (Figure 4B). On day 3, none of the animals in the SAM 10 μg dose group and two out of five in the SAM 30 and 3 μg groups had detectable sgRNA in nasal swabs, compared to four out of five in the control group (Figure 4C), demonstrating significantly decreased viral replication in nasal swabs at the 30 and 10 μg dose levels (p = 0.032(*), 0.008(**) and 0.222(ns), peak viral load for each animal between day 2 and day 10 post challenge compared to control by Mann-Whitney analysis, at 30, 10 and 3 μg, respectively, Supp. Fig. 9). Furthermore, total RNA levels were lower in BAL, oropharyngeal and nasal swabs in all SAM vaccinated animals compared to control animals, with improved viral clearance observed with the 30 and 10 μg of SAM compared with 3 μg SAM (Supp. Fig. 6). Collectively, this data demonstrate protective immunity and efficacy in all SAM vaccinated animals, even at the lowest dose of 3 μg, with the 10 μg and 30 μg dose levels providing comparable and the most effective protection, similar to what was previously described with mRNA vaccines at a dose of 100 μg^8,9,18^.

**Figure 4.**
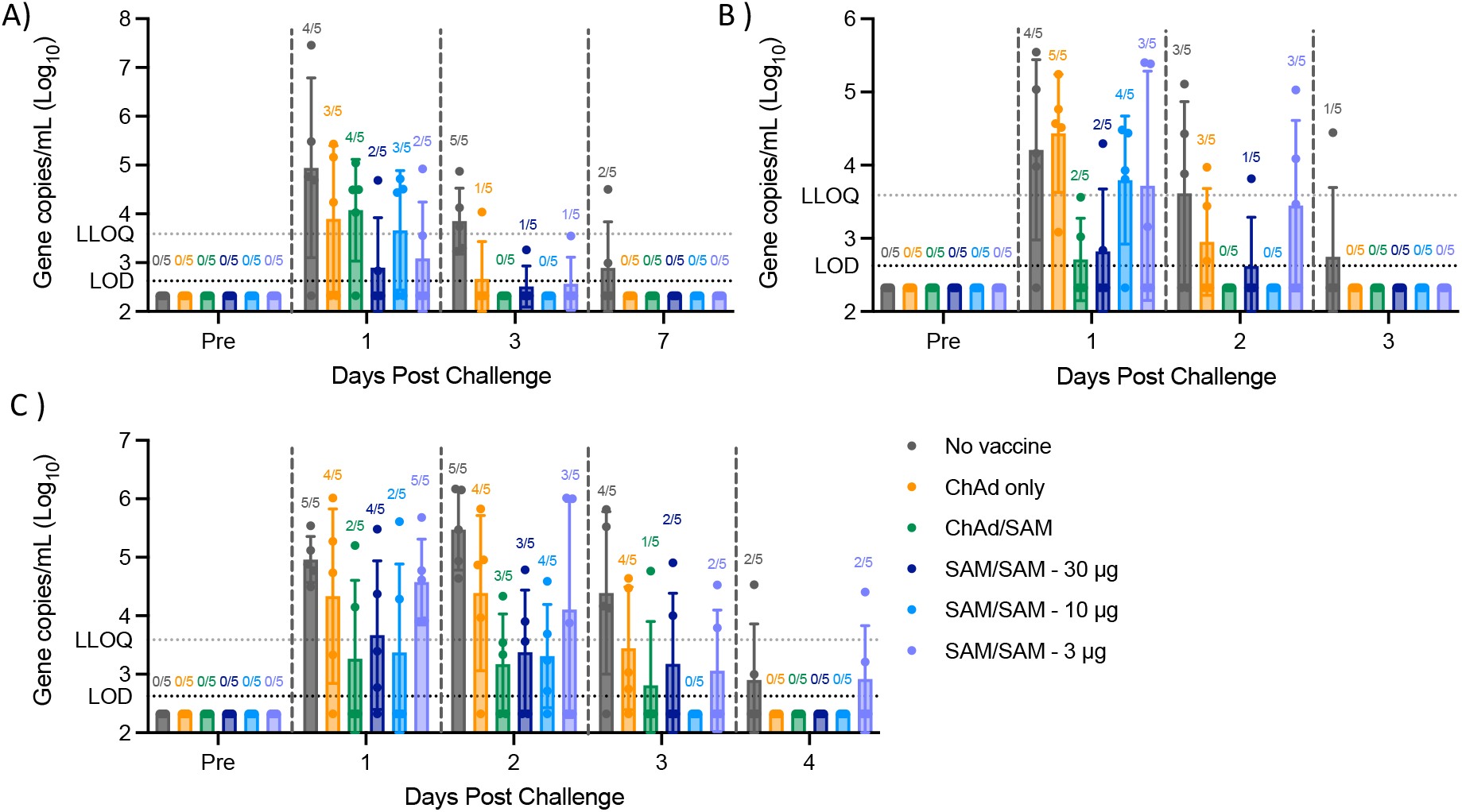
Viral replication is reduced in all vaccinated NHP following SARS-CoV-2 challenge. Subgenomic RNA levels determined by RT-qPCR at specified timepoint post SARS-CoV-2 challenge for each animal in (A) bronchial alveolar lavage (B) oropharyngeal swab or (C) nasal swab. LOD = 422, samples below LOD set to ½ LOD. Geometric mean and SD. Numbers above each bar are numbers of NHP with viral load levels > LOD.

### SAM boost drives strong immune response and protection from infection in ChAd primed rhesus macaques

Since many individuals have already been vaccinated with a SARS-CoV-2 specific adenovirus vaccine, the potency of the SAM vaccine was further explored in a ChAd/SAM heterologous prime/boost vaccine regimen (5×10^11^ VP ChAd prime with 30 μg SAM boost, 6-week interval, Figure 3A), which we have shown to generate potent and durable antigen-specific CD8^+^ T cell responses in ongoing oncology clinical trials (manuscript in preparation). Strong spike S1 IgG antibody titers were observed following a single ChAd immunization and were increased 2.5-fold following SAM boost immunization (GMT 6,847 at 6-weeks post prime and 16,798 2-weeks post boost) to similar titers as those observed following two SAM immunizations at the 30 and 10 μg dose levels (Figure 3B). Strong spike-specific IFNγ, T_H_1 biased, T cell responses were detected 2-weeks after a single immunization of ChAd that were slightly higher than those observed after a single SAM immunization (at any dose), demonstrating the potency of the ChAd vaccine to rapidly drive high titer T cell responses (Figure 3C&D). In addition, T cell responses were increased in 5/5 NHP following the SAM boost (30 μg) immunization to similar levels as those observed following two SAM immunizations (10 or 30 μg), demonstrating the potency of the SAM vaccine as either a boost vector post ChAd prime or as a homologous prime/boost vaccine regimen. Potent neutralizing antibody titers were detected in 10/10 NHP by both PNA and MNA assays following a single ChAd immunization (live virus MNA NT50 GMT 663 4-weeks post prime), again demonstrating the potency of the ChAd vaccine vector. Following SAM (30 μg) boost immunization, titers were increased 2.3-fold (NT50 GMT 2,309 2-weeks post boost), to similar levels as those observed following two doses of 30 or 10 μg SAM (Figure 3E&F), demonstrating comparable potency of the heterologous and homologous prime/boost regimens after two vaccinations even at low doses of the SAM homologous prime/boost vaccination regimen.

Rhesus macaques were challenged with SARS-CoV-2 at 4-weeks following the SAM boost immunization (week 10) and 6-weeks following ChAd prime for the ChAd single immunization group (Figure 3A). A similar increase in neutralizing antibody titers was observed in both groups (Figure 3F). Three days after the challenge, only one of five animals in the ChAd only and zero of five animals in the ChAd/SAM heterologous group had detectable subgenomic RNA in BAL fluid, as compared to five of five animals in the control group. By day 7, none of the animals in any vaccine treated group had detectable sgRNA in the BAL, as compared to 2/5 in the control group (Figure 4A), demonstrating significantly decreased viral replication in the BAL for both a single ChAd immunization and heterologous prime/boost (p = 0.032(*) and p = 0.008(**), respectively, peak viral load for each animal between day 3 and day 10 post challenge compared to control by Mann-Whitney analysis, Supp. Fig. 9). On day 3, one out of five NHP in the heterologous prime/boost group had detectable sgRNA in the nasal swab, compared to four out of five in the ChAd only and control groups (Figure 4C), demonstrating increased protective efficacy with heterologous prime/boost compared to a single ChAd immunization (p = 0.012(*) with ChAd/SAM and p = 0.158(ns) with ChAd only, peak viral load for each animal between day 2 and day 10 post challenge compared to control by Mann-Whitney analysis, Supp. Fig. 9). Collectively, these data demonstrate decreased viral replication and therapeutic benefit provided by the vaccines in all animals, with the 30 μg and 10 μg of SAM homologous prime/boost and the heterologous regimen providing complete protection and the 3 μg SAM homologous prime/boost offering slightly decreased immune response but clearly providing protection to animals challenged with live virus.

## Discussion

Following extensive antigen sequence optimization, we demonstrate that a self-amplifying mRNA vaccine encoding optimized SARS-CoV-2 spike induces potent humoral and cellular immune responses in mice and non-human primates and provides protection from live virus challenge even at low doses. A preliminary SAM-SARS-CoV-2 spike vaccine expressing a full-length spike protein induced strong antigen-specific T cell responses at sub-microgram doses (Supp. Fig. 7) and strong neutralizing antibody responses following a single immunization in mice (Figure 1D), suggesting good potency of this vaccine platform at low doses. The potency of the SAM vaccine platform was compared to an adenovirus vector, a well-studied vaccine platform known to drive robust immune responses, especially T-cell responses, in NHP and humans^19^. The selected adenovirus is based on ChAd68, and both the ChAd68 and SAM vaccine platforms have demonstrated robust neoantigen-specific T cell responses in cancer patients in ongoing clinical trials (manuscript in preparation). Unexpectedly, the SAM vaccine platform expressing our first-generation spike antigen induced significantly higher immune responses in mice compared to the ChAd vaccine, which has not previously been observed with other antigens and thus appeared unique to the spike antigen. This prompted extensive spike sequence optimization efforts, which resulted in selection of a spike sequence that strongly increased antibody titers and T-cell responses in the context of the ChAd vaccine. The SAM vaccine induced immune responses, which were already robust with the original spike sequence, were slightly increased with the optimized spike sequence. This observation highlights the need for careful antigen optimization that is dependent on the antigen and selected vaccine platform. The difference in potency between the two platforms likely reflects differences in the biology of adenovirus-based vaccines compared to SAM vaccines. Adenovirus mRNAs must be transcribed, processed, exported to the cytoplasm, accessible to the ribosome, and stable to produce high levels of protein^20^. As an example, sequence analyses of spike did indeed reveal several splice donor and acceptor sites (data not shown). SAM replication, in contrast, occurs in the cytosol and therefore does not undergo splicing or transport. This might, in part, explain the more potent neutralizing antibody titers induced by the currently authorized mRNA vaccines versus the adenovirus-based vaccine vectors. Nevertheless, human disease severity is still significantly reduced with adenoviral based COVID vaccines^6^, potentially due to the additional induction of robust T-cell responses, which has been demonstrated with this vaccine platform^21^.

We evaluated the SAM vaccine in non-human primates (NHP) to study immune correlates of protection with this alternative self-amplifying mRNA vaccine platform. Strong neutralizing antibody titers were observed following two immunizations of SAM at low doses ranging from 3 to 30 μg. In addition, potent neutralizing antibody titers were generated at the 10 μg dose of SAM even after a single immunization, and increased following a 2^nd^ immunization to levels similar to that observed in rhesus macaques with 10-fold higher doses (100 μg) of authorized SARS-CoV-2 mRNA vaccines^8,9,18^, suggesting that SAM might present a dose sparing vaccine option. Mechanistically, this may be due to the replication of SAM and increased mRNA half-life, which may lead to more potent and potentially more durable immune responses at lower doses compared to non-replicating mRNA vaccines. We are currently evaluating SAM vaccines in humans in ongoing clinical studies for COVID-19. Another COVID-19 SAM vaccine, COVAC1, has been evaluated clinically and vaccine induced immune response was overall lower than the currently authorized mRNA vaccines (Moderna and Pfizer/BioNTech), with only a 61% seroconversion rate at the 10 μg dose^22^, compared to 100% with both the Pfizer/BioNTech mRNA vaccine at doses ranging from 10 – 30 μg^23^ and the Moderna mRNA vaccine at 50 and 100 μg^24^. This decrease in potency may have resulted from activation of innate immune pathways that can partly restrict SAM replication and antigen expression and vary across individuals, which may be triggered by both the LNP and/or the self-amplifying RNA itself. Another SAM vaccine platform that uses an alternative non-prefusion stabilized spike sequence and a different LNP formulation, ARCT-021, demonstrated 100% seroconversion clinically at low doses (3 – 7.5 μg), underscoring the impact of the spike sequence and structural and chemical differences of SAM vaccine platforms on vaccine potency^25^. However, ARCT-021 induced low neutralizing antibody titers which were below the GMT titer of convalescent sera, demonstrating decreased immunogenicity compared to the Pfizer/BioNTech and Moderna mRNA vaccines. Notably, as shown herein, our SAM vaccine demonstrates increased immunogenicity in mice compared to both the COVAC1 and ARCT-021 SAM vaccine platforms (note no published data in non-human primates available for these vaccines) (Figure 1D, Supp. Fig. 2G, 7), which might be driven by use of different formulations as well as structural differences in both the spike sequence, such as codon optimization and prefusion stabilizing modifications, and differences in the SAM backbone including the 5’ untranslated sequence. During the preclinical development of our SAM SARS-CoV-2 vaccine, we were able to demonstrate that each of these components clearly impact the vaccine induced immune response.

One difference between the authorized mRNA vaccines and the SAM vaccines in development is the use of modified nucleosides in the mRNA vaccines, which have been shown to reduce innate immune response post vaccination and may lead to increases in the vaccine induced adaptive immune response. The lack of modified nucleotide usage in the mRNA vaccine developed by CureVac, which demonstrated lower clinical efficacy compared to the Moderna and Pfizer/BioNTech vaccines^26^, supports this concept. These observations suggest that controlling the innate immune response early during vaccination results in an increased vaccine induced immune response. We have studied the induction of innate immune response to SAM carefully during the preclinical development of our SAM vaccine and optimized various components in the vaccine to decrease the activation of innate immune pathways. Our data in NHP suggest that our SAM SARS-CoV-2 vaccine does not lead to strong systemic innate immune activation at the two lower doses of 3 and 10 μg, demonstrated by the lack of increased IFNα serum levels following vaccination, as was observed with the 30 μg dose in this study (Supp. Fig. 8) and in previous NHP studies at higher doses of SAM (data not shown). Induction of innate immunity and its impact on vaccine potency might differ between mRNA and SAM vaccines, due to the replication of the SAM vaccine, which is an area of intense investigation.

Spike-specific T_H_1 biased T cell responses were observed in SAM immunized NHP at doses ranging from 3 μg to 30 μg, although with high heterogeneity across individual animals, which may reflect the diversity of MAMU haplotypes and the small group size, as animals were not randomized based on their MAMU haplotype. Accumulating evidence has demonstrated the importance of cellular immunity in clearance of virus infected cells, protection from SARS-CoV-2 infection, and reduction of disease severity^27,28^. CD8 depletion studies in NHP have demonstrated that SARS-CoV-2 virus specific CD8^+^ T cells contribute to protection from SARS-CoV-2, particularly in the context of waning antibody titers^16^. Similarly, strong T cell responses are associated with improved survival in COVID-19^+^ patients with hematologic cancer, who have B cell deficiency and therefore mount weak humoral responses post immunization with the currently approved COVID vaccines^29^. T cells may also contribute to broader vaccine protection against SARS-CoV-2 variants of concern, despite the reduced neutralization provided by the spike antibodies^30,31^. Furthermore, T-cell memory may be more durable than B-cell memory and thus induction of broad T-cell memory against SARS-CoV-2, in addition to neutralizing antibodies, may provide longer and more potent protection against future coronavirus variants and should be considered in the development of panCOVID vaccines^32^.

Following challenge with SARS-CoV-2, a rapid increase in neutralizing antibody response was observed in all vaccinated animals, demonstrating an anamnestic response, which potentially contributed to the observed rapid clearance of viral load and suppression of viral replication. Notably, viral load in unvaccinated control animals following challenge was higher in this study when compared to published NHP challenge studies evaluating other vaccine regimens, potentially due to use of a more potent live virus lot, the challenge technique used, or assay variability in assessing viral load^8,9^. Effective protection as measured by viral replication was observed with all vaccine regimens, with homologous SAM prime/boost demonstrating protection and decreased viral load at doses as low as 3 μg.

We have demonstrated that homologous SAM prime/boost drives potent humoral and cellular immune responses and provides protection from SARS-CoV-2 infection at low doses in NHP. The SAM vaccine (encoding the Beta (501Y.V2) spike variant in addition to T-cell epitope sequences targeting regions of the virus outside of spike) will be evaluated in a Coalition for Epidemic Preparedness Innovations (CEPI) sponsored clinical trial in South Africa.

In this study, we have further demonstrated that SAM can provide a powerful boost following a ChAd prime vaccination, driving potent T-cell and antibody responses. The heterologous prime/boost concept has recently been evaluated clinically using the currently approved Ad and mRNA vaccines^11,12^. In one study, it was demonstrated that mRNA heterologous boosts led to comparable or increased antibody responses, compared to homologous boosts^11^. Another study demonstrated that an mRNA boost following the AstraZeneca ChAdOx1 prime results in improved spike CD8^+^ T-cell responses compared to two mRNA immunizations^12^. This “mix-and-match” approach has been approved by the FDA. While a single vaccine platform is preferred for simplicity of manufacturing and rollout during a pandemic, a heterologous ChAd prime and SAM boost regimen may drive more rapid, stronger and more consistent T cell responses, and therefore, may offer an attractive and potent option to a subset of individuals who, for example, have weakened immune systems or are immune compromised, such as the elderly or cancer patients on chemotherapy. We have shown that our ChAd/SAM prime/boost vaccine regimen drives robust neoantigen-specific T cell responses in cancer patients, even in patients who received concurrent chemotherapy, demonstrating the potency of this heterologous prime/boost approach^33^ (Clinical trials NCT03639714, NCT03953235, manuscript in preparation). The heterologous prime/boost concept is currently being evaluated in a clinical trial in the UK, utilizing our SARS-CoV-2 SAM vaccine as a boost, at 10 or 30 μg, in individuals >60 years old who have received the AstraZeneca ChAdOx1vaccine as a primary series and in a DMID sponsored study in the US in individuals who have been primed with the J&J Ad26 based vaccine (NCT04776317).

In summary, the SAM vector is an emerging and potent vaccine platform that drives robust T-cell and neutralizing antibody responses at low doses, protects NHP from SARS-CoV-2 and is currently being tested in humans in both homologous and heterologous prime/boost regimens. It is rapidly adaptable, simple to manufacture, and quick to release, offering an additional attractive vaccine platform in the fight against the ongoing SARS-CoV-2 pandemic and current and emerging infectious pathogens, as well as cancer.

## Acknowledgements

We thank Nexelis for running the pseudovirus neutralization assay on mouse sera and their support in transferring the assay for assessing non-human primate sera. We thank Battelle Memorial Institute for execution of the non-human primate challenge study and performing the live virus microneutralization and RT-PCR assays.

## Author Contributions

CDS, LG, SJH, and KJ designed vaccine constructs. CDS, SJH, JAE, RLV, LG, AA and MF produced the vaccines and performed in vitro evaluation. ARR, AS, AVA, ME, MAK and GB designed and executed mouse studies and performed and analyzed serum IgG and ELISpot assays. ARR and GS designed NHP study. GB performed and analyzed NHP ELISpot assays and cytokine MSD. MAK performed and analyzed NHP PNA assays and ME performed and analyzed NHP IgG assays. GS oversaw performance and analysis of MNA and RT-PCR assays. ARR, SJH, CDS and KJ wrote the manuscript.

## Methods

### Codon Optimization

Spike nucleotide sequences based on the original Wuhan-Hu-1 strain (MN908947) with the D614G mutation were designed for maximum expression in Homo Sapiens using three different codon optimization tools. Spike-V1 was generated using the IDT (Coralville IA) optimization tool, V2-V7 were codon optimized using the COOL optimization algorithm^34^ and V8 was optimized using an SGI-DNA (La Jolla, CA) codon optimization tool. Each codon optimized sequence was ordered as double stranded DNA gBlocks (IDT) with at least a 30 bp overlap with sequences flanking the ChAd68 cloning site.

### Vector generation

The ChAd68 nucleotide sequence was based on the wild-type sequence obtained by MiSeq (Ilumina sequencing) of virus obtained from the ATCC (VR-594). The sequence of a E1 (578-3404 bp)/E3 deleted virus (2,125-31,825 bp) was assembled into pUC19 from VR-594-derived and synthetic (SGI-DNA) fragments. An E4 deletion between E4ORF2-4 was introduced by PCR. A CMV promoter/enhancer with an SV40 polyA was introduced into the E1 region and the spike gBlock sequences introduced by Gibson assembly (Codexis) and transformed into Stbl4 (Thermo Fisher) cells. Error free clones were selected by PCR and sequencing and plasmid DNA prepared at the Maxi-prep scale (Machery-Nagel). Furin and proline spike mutations, 2P or 6P, were introduced into the spike protein by overlapping PCR extension using primers to introduce the specific mutations.

The pA68-E4d-Spike plasmids were linearized, purified using a Nucleospin kit (Machery-Nagel) and transfected into 2 mL of 293F cells (0.5 mL/mL) using TransIT-Lenti (Mirus bio). The virus was amplified, harvested and re-infected into 30 mL of 293F cells for 48-72h. Cells and media were harvested and used to infect 400 mL of 293F cells. Cells were harvested after 48 hours and lysed by a freeze/thaw step (−80°C/37°C) in 10mM Tris pH 8.0/0.1% Triton-X100, and then purified by two successive rounds of CsCl gradient centrifugation. Virus bands were purified and dialyzed 3x into 1X ARM buffer (10 mM Tris pH 8.0, 25 mM NaCl, 2.5% glycerol). Viral particle concentration was determined by the Absorbance 260 nm method post lysis in 0.1% SDS and the infectious unit (IU) titer was determined by immunostaining as described previously^35^.

Spike sequences were PCR amplified and cloned into PacI/BstBI sites of a pUC02-VEE vector. Capped SAM was synthesized in vitro using HiScribe T7 Quick High Yield RNA Synthesis Kit (New England Biolabs) and purified using a RNeasy Maxi Kit (Qiagen) according to the manufacturer’s protocol. SAM was subsequently encapsulated in a lipid nanoparticle (LNP) using a self-assembly process in which an aqueous solution of SAM is rapidly mixed with a lipid mixture in ethanol^36^. RNA encapsulation efficiency was measured using Ribogreen RNA quantitation reagent (Thermo Fisher) and confirmed to be >95% in all batches analyzed. SAM-LNP was formulated into a buffer containing 5 mM Tris (pH 8.0), 10% sucrose, 10% maltose.

### Western analysis

HEK293F cells seeded at 5e5 cells/mL were infected with an MOI of 1 IU/cell. Cells were incubated between 40-72h before harvesting by centrifugation. Cells were washed in 1xPBS and resuspended in 1XSDS/PAGE buffer with 2.5% β-mercaptoethanol, then sheered through a 27g needle, prior to incubation at >90°C for 5 min. Samples (10μL) were separated on a 4-20% SDS-PAGE gel and blotted onto a PVDF membrane using a Trans-Blot Turbo transfer system (BioRad). Membranes were blocked for 1h in 5% skim milk in TBST and then probed with an anti-S2 mouse monoclonal antibody (GeneTex) at a 1:1000 dilution for 2h-overnight. Membranes were washed (0.05% Tween 20 in 1X TBS) and then probed with a Rabbit anti-Mouse HRP antibody (Bethyl labs) for 1h before washing and detection with a SuperSignal West Femto Maximum Sensitivity Substrate (Pierce). A parallel blot using a mouse anti-actin antibody (Thermo Fisher) was used to ensure equivalent protein amounts per well.

### Mouse studies

Mouse studies were conducted at Murigenics (Vallejo, CA) under IACUC approved protocols. Female Balb/c mice (Envigo), 6–8 weeks old were used for all studies. All immunizations were bilateral intramuscular to the tibialis anterior, 2 injections of 50 μL each. Mouse spleens were harvested 12–14 days following immunization. Spleens were suspended in RPMI complete (RPMI + 10% FBS) and dissociated using the gentleMACS Dissociator (Milltenyi Biotec). Dissociated cells were filtered using a 40 μm strainer and red blood cells were lysed with ACK lysing buffer (150 mM NH_4_Cl, 10 mM KHCO_3_, 0.1 mM EDTA). Following lysis, cells were filtered with a 30 μm strainer and resuspended in RPMI complete. At various timepoints post immunization, 200 μL of blood was drawn. Blood was centrifuged at 1000g for 10 minutes at room temperature. Serum was collected and frozen at –80°C.

### Intracellular Cytokine Staining

Freshly isolated splenocytes were resuspended at a density of 5×10^6^ cells/mL in complete RPMI and following an overnight rest at 4°C, 1×10^6^ cells per well were distributed into v-bottom 96-well plates. Cells were pelleted and resuspended in 100 μL of complete RPMI containing an overlapping peptide pool containing 316 peptides (each 15 amino acids in length, 11 amino acid overlap) spanning the SARS-CoV-2 spike antigen, at a final concentration of 0.5 μg/mL per peptide (Genscript). A second well with DMSO only was used as a negative control for each sample. After 1 hour of incubation at 37ºC, Brefeldin A (Biolegend) was added to a final concentration of 5 μg/mL and cells were incubated for an additional 4 hours. Following stimulation, cells were washed with PBS and stained with fixable viability dye (eBioscience). Extracellular staining was performed in FACS buffer (PBS + 2% FBS + 2mM EDTA) with the following antibodies: CD4 (GK1.5, Biolegend), CD8 (53-6.7, BD). Cells were then washed, fixed and permeabilized with the eBiosciences Fixation/Permeabilization Solution Kit. Intracellular staining was then performed in permeabilization buffer with the following antibodies: IFNγ (XMG21.2, Invitrogen), TNFα (MP6-XT22, eBiosciences), IL2 (JES6-5H4, eBiosciences), IL4 (11B11, Biolegend), IL10 (JES5-16E3, Biolegend). Samples were collected on a Cytoflex LX (Beckman Coulter). Analysis of flow cytometry data was performed using FlowJo software.

### Non-human primate studies

Study was conducted in compliance with all relevant local, state and federal regulations and were approved by the Battelle Institutional Animal Care and Use Committee (IACUC). 30 Chinese-origin male and female rhesus macaques (*M. mulatta)* >2.5 years old were housed at Battelle (Columbus, Ohio). NHP were vaccinated with either ChAd-Spike(V2)-F2P (Group 1 - 5×10^11^ VP, study day 0; Group 2 – 5×10^11^ VP, study day 28), SAM-Spike(V2)-F2P (Group 1 - 30 μg, study day 42; Group 3 – 30 μg, study days 14 and 42; Group 4 - 10 μg, study days 14 and 42; Group 5 - 3 μg, study days 14 and 42), or PBS (Group 6, study days 0 and 42). All injections were bilateral intramuscular, 0.5 mL per leg (1 mL total) to the thigh. Note that immunizations were staggered to enable challenge immunization on study day 70 for all groups. Each group was divided into two sub-groups and study initiated one week apart, to enable two staggered challenge groups. Animals were challenged with SARS-CoV-2 (strain USA-WA1/2020) via the intratracheal (0.5 mL) and intranasal (0.25 mL per nostril) routes with a target dose of ∼1.6 × 10^6^ PFU (undiluted). Serum was collected using serum separator tubes and whole blood collected in cell preparation tubes and peripheral blood mononuclear cells isolated and cryopreserved in CryoStorCS solution (Sigma Aldrich).

### ELISpot assays

IFNγ ELISpot assays were performed using pre-coated 96-well plates (Mabtech, Monkey IFNγ ELISPOT PLUS, ALP or Mouse IFNγ ELISPOT PLUS, ALP) following manufacturer’s protocol. For NHP, frozen PBMCs were thawed at 37°C and then rested overnight in RPMI + 10% FBS. 1×10^5^ PBMCs were plated per well in triplicate with a single overlapping peptide pool spanning spike from the N to C terminus (GenScript, 15 amino acid length, 11 amino acid overlap, 314 peptides total) at final concentration of 1 ug/mL per peptide and incubated overnight at 37°C in RPMI + 10% FBS. For mouse studies, freshly isolated splenocytes were stimulated overnight with either two (∼120 peptides/each) or eight different overlapping peptide pools (36 – 40 peptides each) spanning the SARS-CoV-2 spike antigen, at a final concentration of 1 μg/mL per peptide (GenScript). Splenocytes were plated in duplicate at 1×10^5^ cells per well and 2.5×10^4^ cells per well (mixed with 7.5×10^4^ naïve cells) for each stimulus. DMSO only was used as a negative control for each sample. Plates were washed with PBS and then incubated with anti-monkey or anti-mouse IFNγ mAb biotin (Mabtech) for two hours, followed by an additional wash and incubation with Streptavidin-ALP (Mabtech) for one hour. After final wash, plates were incubated for ten minutes with BCIP/NBT (Mabtech) to develop the immunospots. Wells were imaged and spots enumerated using AID reader (Autoimmun Diagnostika). Samples with replicate well variability (Variability = Variance/(median + 1) greater than 10 and median greater than 10 were excluded^37^. Spot values were adjusted based on the well saturation according to the formula: AdjustedSpots = RawSpots + 2*(RawSpots*Saturation/(100-Saturation)^38^. Each sample was background corrected by subtracting the average value of the negative control wells. Data was normalized to spot forming units (SFU) per 1×10^6^ cells by multiplying the corrected spot number by 1×10^6^/cell number plated. Data processing was performed using the R programming language and graphed using GraphPad Prism. Statistical analysis was performed in GraphPad Prism.

### Pseudovirus neutralization assay

Mouse and human convalescent serum samples (courtesy of Helen Chu, University of Washington) were assessed by Nexelis (Laval, Quebec). NHP serum samples were assessed by Gritstone bio (Emeryville, CA) using the same pseudovirus, controls, reagents and protocol. Pseudotyped virus particles were made using a genetically modified Vesicular Stomatitis Virus from which the glycoprotein G was removed (VSVΔG). The VSVΔG virus was transduced in HEK293T cells previously transfected with the spike glycoprotein of the SARS-CoV-2 coronavirus (Wuhan strain) for which the last 19 amino acids of the cytoplasmic tail were removed (ΔCT). The generated pseudovirus particles (VSVΔG – Spike ΔCT) contain a luciferase reporter which can be quantified in relative luminescence units (RLU). Heat-inactivated serum samples were serially diluted (7-serial 2-fold dilution) in a 96-well plate and a pre-determined amount of pseudotyped virus (corresponding to between approximately 75,000 and 300,000 RLU/well) was applied to the plate and incubated with serum/plasma to allow binding of the neutralization antibodies to the pseudotyped virus.

After the incubation of the serum/plasma-pseudotyped virus complex, the serum/plasma-pseudotyped virus complex was transferred to the plate containing Vero E6 cells (ATCC). Test plates were incubated at 37°C with 5% CO2 overnight. Luciferase substrate was added to the plates which were then read using a plate reader detecting luminescence. The intensity of the light being emitted is inversely proportional to the amount of anti-SARS-CoV-2 neutralizing Spike antibodies bound to the VSVΔG – Spike ΔCT particles.

Each microplate was read using a luminescence microplate reader (SpectraMax). The dilution of serum required to achieve 50% neutralization (NT50) when compared to a non-neutralized pseudoparticle control was calculated for each sample dilution and the NT50 is interpolated from a linear regression using the two dilutions flanking the 50% neutralization.

### Microneutralization assay

The microneutralization assay was performed at Battelle Memorial Institute (Columbus, Ohio) to assess the neutralizing antibody titer in serum samples collected from NHPs following vaccination and post-challenge. The virus neutralization titer is expressed as the reciprocal value of the highest dilution of the serum which still inhibited virus replication. All serum samples were analyzed in duplicate.

Briefly, 2-fold serial dilutions of heat-inactivated serum sample were pre-incubated with the virus for 60 minutes at 37°C. The virus/serum mixture was then added to a 90 to 100% confluent monolayer of Vero E6 cells (BEI, Cat. No. NR-596) in 96-well plates and incubated for two days at 37°C with 5% CO_2_. Following incubation, the inoculum was removed, and monolayers were incubated for 30 minutes in 80% cold acetone to allow cell fixation. Plates were incubated with anti-nucleocapsid protein primary antibody cocktail (clones HM1056 and HM1057) (EastCoast Bio, North Berwick, ME) for 60 minutes at 37°C (Battelle Memorial Institute, Patent Number 63/041,551 Pending, 2020). The plates were washed and the secondary antibody (goat anti-mouse IgG Horse Radish Peroxidase (HRP) conjugate; Fitzgerald, North Acton, MA) was added to the wells, and the plates were incubated for 60 minutes at 37°C^1^. After the plates were washed, the substrate was added, and the plates were incubated at 37°C. Stop solution was added, and the plates were read for optical density at 405 nm wavelength. Neutralizing activity is defined as at least 50% reduction in signal from the virus only (VC) wells relative to cells control (CC) wells following the formula [(average VC – average CC)/2] + average CC. The median neutralizing titer (MN_50_) was calculated using Spearman-Kärber analysis method^39^.

### S1 IgG ELISA

96-well QuickPlex plates (Meso Scale Discovery, Rockville, MD) were coated with 50 μL of 1 μg/mL SARS-CoV-2 S1 (ACROBiosystems, Newark, DE), diluted in DPBS (Corning, Corning, NY), and incubated at 4°C overnight. Wells were washed three times with agitation using 250 μL of PBS + 0.05% Tween-20 (Teknova, Hollister, CA) and plates blocked with 150 μL Superblock PBS (Thermo Fisher Scientific, Waltham, MA) for 1 hour at room temperature on an orbital shaker. Test sera was diluted at appropriate series in 10% species-matched serum (Innovative Research, Novi, MI) and tested in single wells on each plate. Wells were washed and 50 μL of the diluted samples were added to wells and incubated for 1 hour at room temperature on an orbital shaker. Wells were washed and incubated with 25 μL of 1 μg/mL SULFO-TAG labeled anti-species antibody (MSD), diluted in DPBS + 1% BSA (Sigma-Aldrich, St. Louis, MO), for 1 hour at room temperature on an orbital shaker. Wells were washed and 150 μL Read Buffer T (MSD) added. Plates were run immediately using the QPlex SQ 120 (MSD) ECL plate reader. For each sample, the endpoint titer was calculated as the reciprocal serum dilution at 2-fold the average background value for the plate, interpolated by a linear regression using the two highest dilutions that gave values greater than 2-fold the average background.

### IFN**α**2**α** ELISA

NHP serum samples were analyzed for levels of IFNα2α using a MSD U-PLEX Biomarker assay (catalog number K15068L-2), according to the manufacturer’s instructions. Analyte concentration (pg/mL) was calculated using serial dilutions of known standards. Each animal and timepoint was run in technical duplicates.

### Viral RNA RT-qPCR

#### Nucleocapsid Protein (N1) Genomic Analysis

Briefly, RNA was isolated using the Indispin QIAcube HT Pathogen Kit (Indical Bioscience, Germany) on the QIAcube HT instrument (Qiagen, Germany). The isolated RNA was then evaluated in RT-qPCR using the TaqMan Fast Virus 1-step Master Mix (Thermo Fisher Scientific) on a QuantStudio Flex 6 Real-Time PCR System (Applied Biosystems; Foster City, CA). The primers and probe were specific to the SARS-CoV-2 nucleocapsid gene, corresponding to the N1 sequences from the Centers for Disease Control and Prevention (CDC) 2019-Novel Coronavirus (2019-nCoV) Real-Time RT-PCR Diagnostic Panel (https://www.cdc.gov/coronavirus/2019-ncov/lab/rt-pcr-panel-primer-probes.html) except that the probe quencher was modified to Non-Fluorescent Quencher-Minor Groove Binder (NFQ-MGB) (Thermo Fisher Scientific). A standard curve comprised of synthetic RNA containing the target sequence from SARS-CoV-2 isolate WA1 sequence (GenBank Accession Number MN985325.1) (Bio-Synthesis, Inc.; Lewisville, TX) was included on each PCR plate for absolute quantitation of SARS-CoV-2 RNA copies in each sample. Thermocycling conditions were as follows: Stage 1 - 50°C for 5 min for one cycle; Stage 2 - 95°C for 20 sec for one cycle; Stage 3 - 95°C for 3 sec and 60°C for 30 sec for 40 cycles. Data analysis was performed using the QuantStudio 6 software-generated values (total copies per well of each sample) and additional calculations to determine SARS-CoV-2 N1 copies per mL of fluid.

#### Envelope Protein (E) Subgenomic Analysis

Following isolation and evaluation using the N1 genomic assay, the isolated RNA was then evaluated as described above using primers and probes specific to the SARS-CoV-2 E gene based on previously described sequences^40^, and the reverse primer and probe sequences previously described^41^ (Integrated DNA Technologies, Iowa). A standard curve comprised of synthetic RNA containing the target sequence from SARS-CoV-2 isolate WA1 sequence (GenBank Accession Number MN985325.1) (Bio-Synthesis, Inc.; Lewisville, TX) was included on each PCR plate for absolute quantitation of SARS-CoV-2 copies in each sample. Thermocycling conditions were as follows: Stage 1 - 50°C for 5 min for one cycle; Stage 2 - 95°C for 20 sec for one cycle; Stage 3 - 95°C for 3 sec and 60°C for 30 sec for 40 cycles. Data analysis was performed using the QuantStudio 6 software-generated values (total copies per well of each sample) and additional calculations to determine SARS-CoV-2 E gene subgenomic (Esg) RNA copies per mL of fluid.

---

**Supplemental Figure 1.**
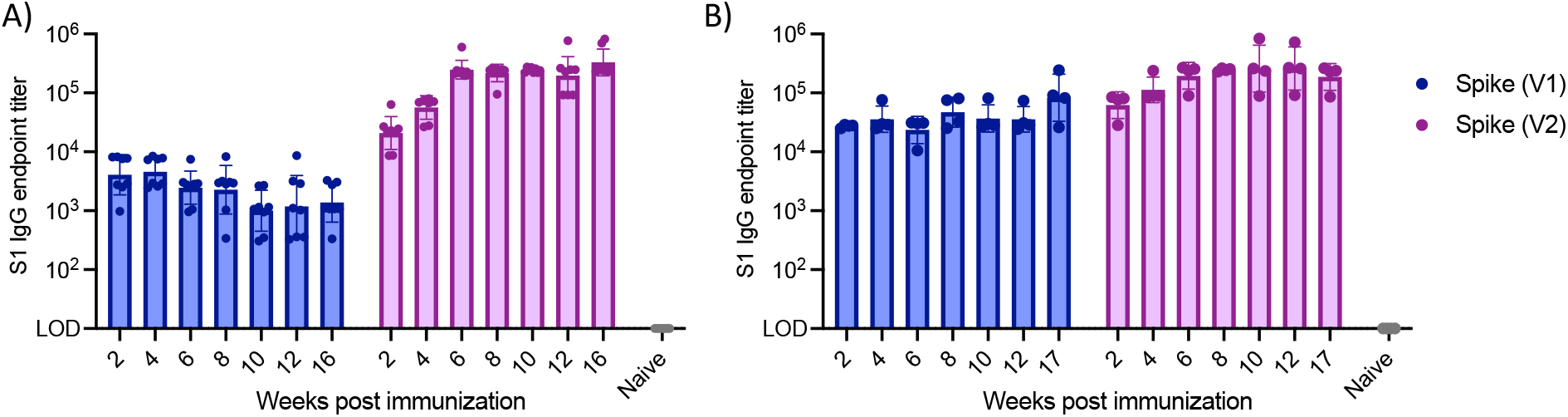
Serum spike S1 IgG antibody binding titers in Balb/c mice assessed by MSD ELISA at the specified timepoint post immunization with either (A) ChAd (1×10^11^ VP) or (B) SAM (10 μg) encoding either the Spike V1 or V2 sequence. Naïve samples all below LOD (LOD = 10). Geomean and geometric SD.

**Supplemental Figure 2.**
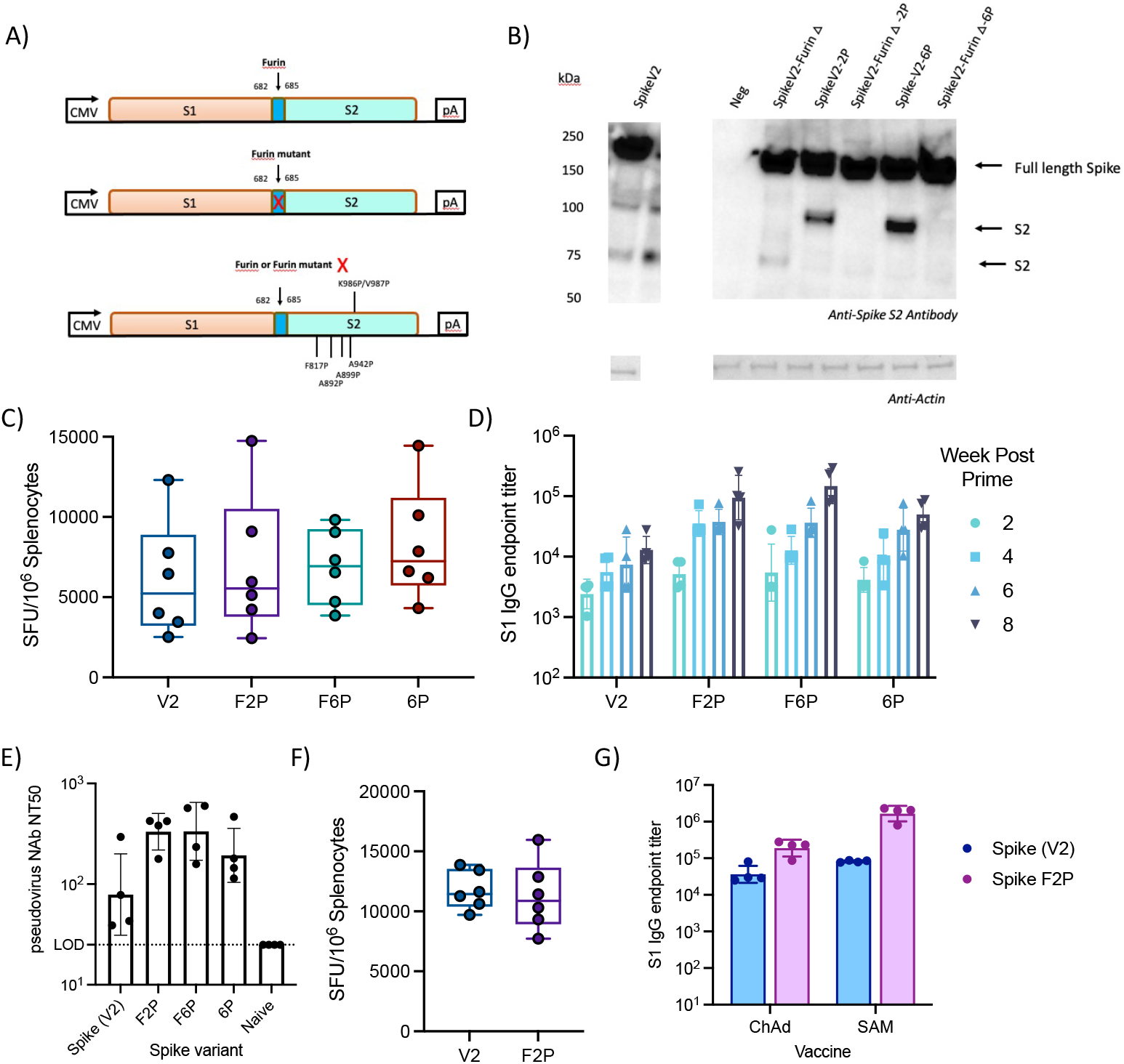
Addition of Furin mutation and proline substitutions increases neutralizing antibody titers in mice. (A) Schematic of ChAd constructs expressing a codon optimized spike V2 either as a wild-type protein or with mutations in the Furin cleavage site and/or with proline stabilization mutations: WT-Spike, wild-type spike; Furin△Spike, Furin or Furin mutated spike with 2-proline, K986P/V987P (2P) or 6-proline mutations F817P, A892P, A899P, A942P, K986P & V987P (6P). (B) Western blot of lysates analyzed 72h post infection (MOI 1) with ChAd-SpikeV2 constructs or with a negative control ChAd (Neg) expressing a non-spike protein, using an anti-S2 antibody. An anti-Actin antibody was used as loading control. (C) Spike specific T cell response in Balb/c mice splenocytes 2 weeks post immunization with ChAd vaccine encoding the specified spike variant (1×10^10^ VP), IFNγ ELISpot, sum of eight spike peptide pools. Median, IQR, range. (D) Serum spike S1 IgG antibody binding titers in Balb/c mice assessed by MSD ELISA at the specified timepoint post immunization with ChAd vaccine encoding the specified spike variant (1×10^10^ VP). (E) Serum pseudovirus neutralization titers (50% inhibition) in Balb/c mice 4 weeks post immunization with ChAd vaccine (1×10^10^ VP) encoding the specified spike variant. (F) Spike specific T cell response in Balb/c mice splenocytes 2 weeks post immunization with SAM (10 μg) vaccine encoding the specified spike variant, IFNγ ELISpot, sum of two spike peptide pools. Median, IQR, range. (G) Serum spike S1 IgG antibody binding titers in Balb/c mice assessed by MSD ELISA at 4 weeks post immunization with ChAd (1×10^11^ VP) or SAM (10 μg) vaccine encoding the specified spike variant. Geomean and geometric SD.

**Supplemental Figure 3.**
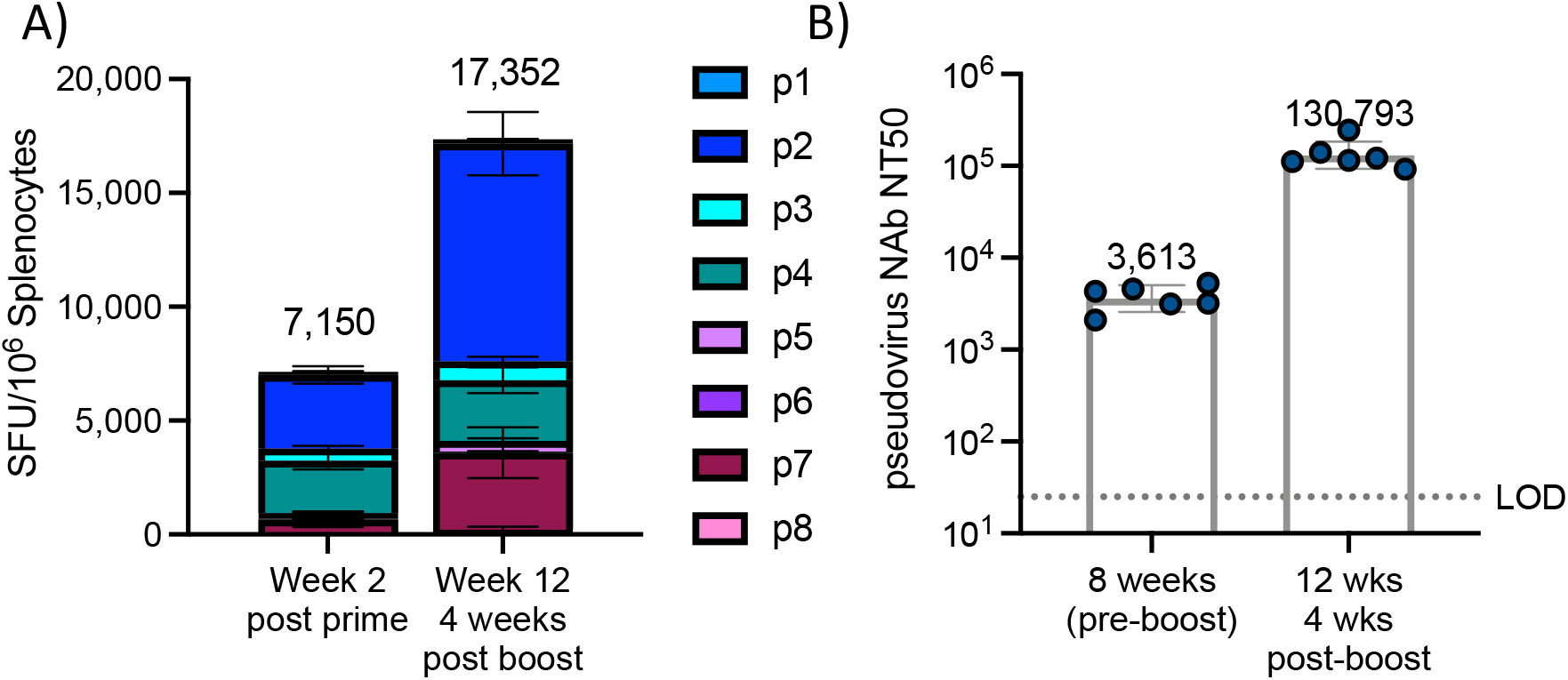
Heterologous prime/boost in Balb/c mice. Mice immunized with ChAd (1×10^11^ VP) prime, followed by SAM (10 μg) boost 8 weeks post prime. N = 6/group. (A) IFNγ ELISpot at the specified timepoint, following stimulation with eight peptide pools spanning spike antigen. Mean ± SEM for each peptide pool. (B) Pseudovirus neutralizing titer (NT50) at the specified timepoint. Geometric mean and geometric SD. Sera from Naïve mice were below LOD of 25.

**Supplemental Figure 4.**
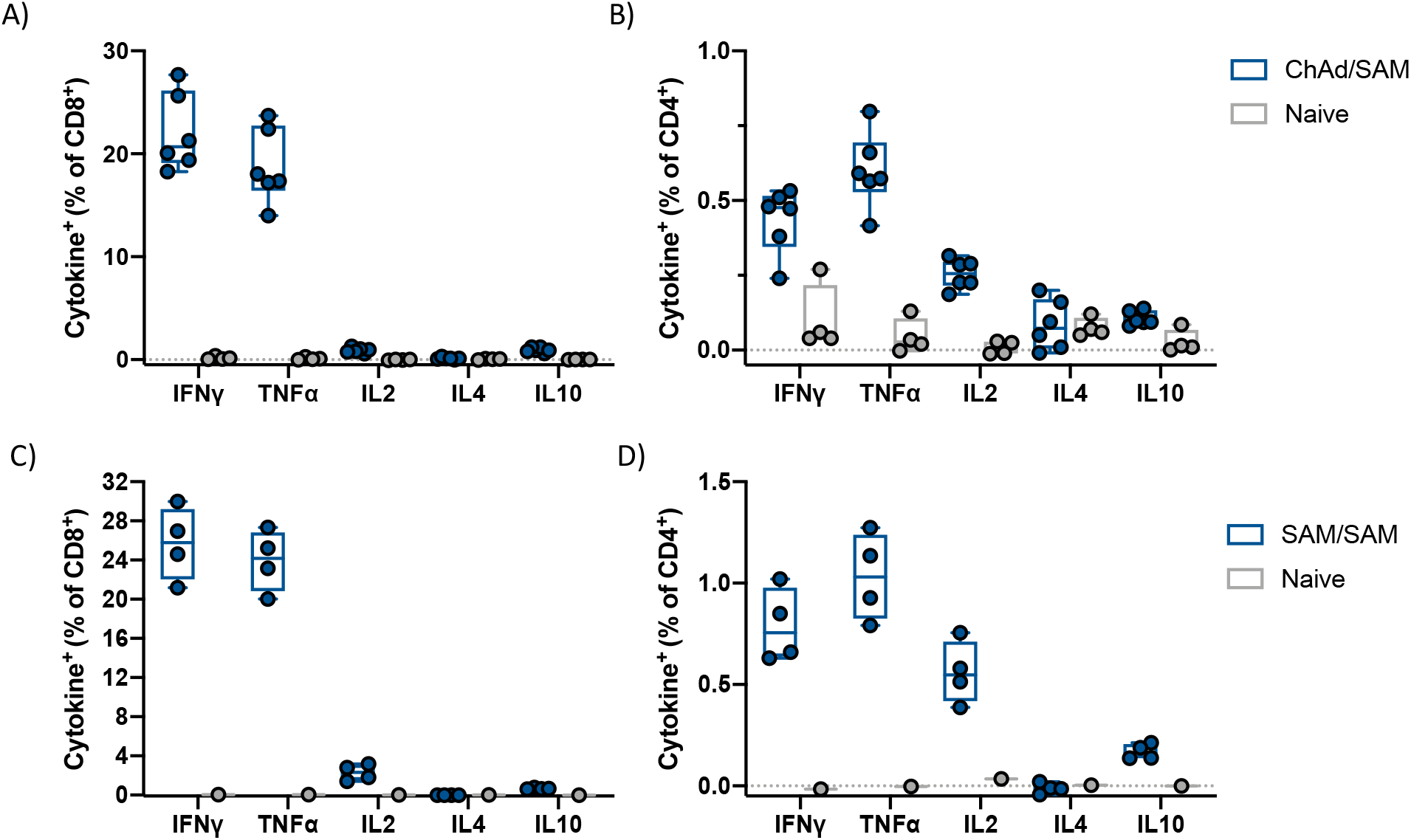
Vaccine induced T cell response is T_H_1 biased. Intracellular cytokine staining in splenocytes of Balb/c mice following stimulation with overlapping peptide pool spanning spike. Background corrected, box and whiskers represent median, IQR, and range. (A-B) Mice immunized with 1×10^11^ VP ChAd-Spike(V2)-F2P and boosted with SAM-Spike(V2)-F2P (10 μg) at 8 weeks. T cell response assessed at week 12, 4 weeks post boost (n=6) compared to unvaccinated mice (n=4). (C-D) Mice immunized with SAM-Spike(V2)-F2P (10 μg) at 0 and 8 weeks. T cell response assessed at week 10, 2 weeks post boost (n=4) compared to unvaccinated mice (n=1). (A&C) Cytokine positive out of CD8+ T cells (%) (B&D) Cytokine positive out of CD4+ T cells (%).

**Supplemental Figure 5.**
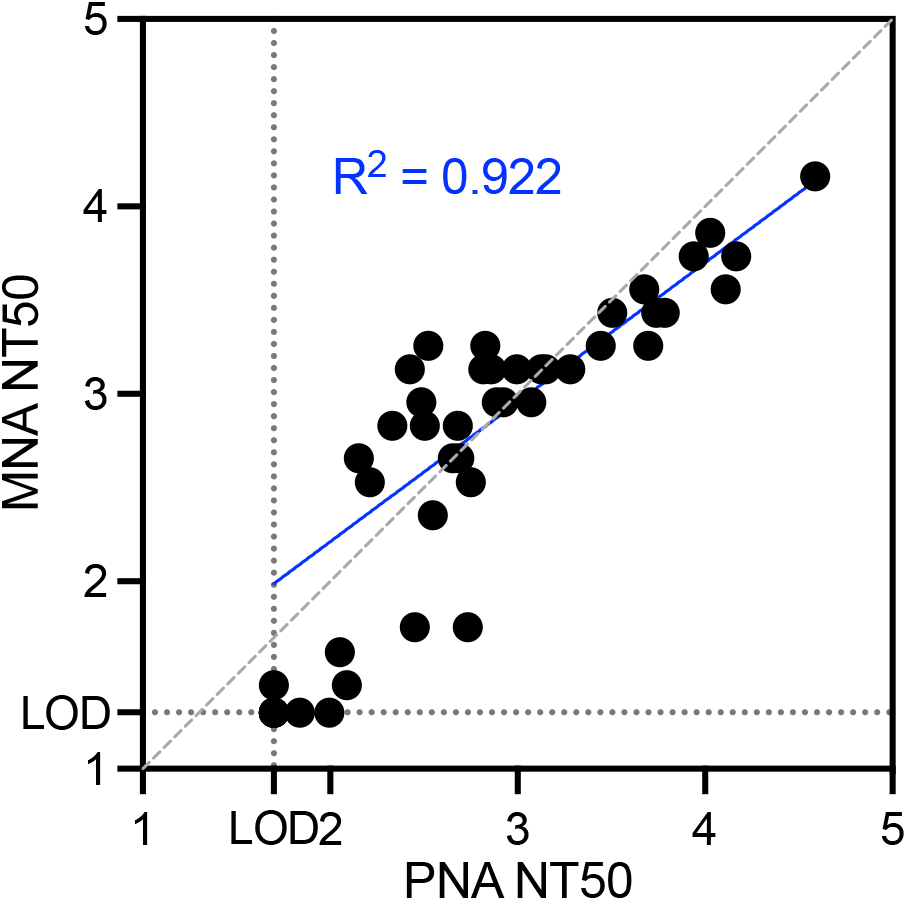
Scatterplot comparing NT50 titers measured by pseudovirus neutralization assay (PNA) and live virus microneutralization assay (MNA) for all samples that were assessed by both assays (excluding baseline samples). Dashed gray line represents unity. LOD for MNA assay = 20, LOD for PNA assay = 50. Blue line represents non-linear regression analysis best fit.

**Supplemental Figure 6.**
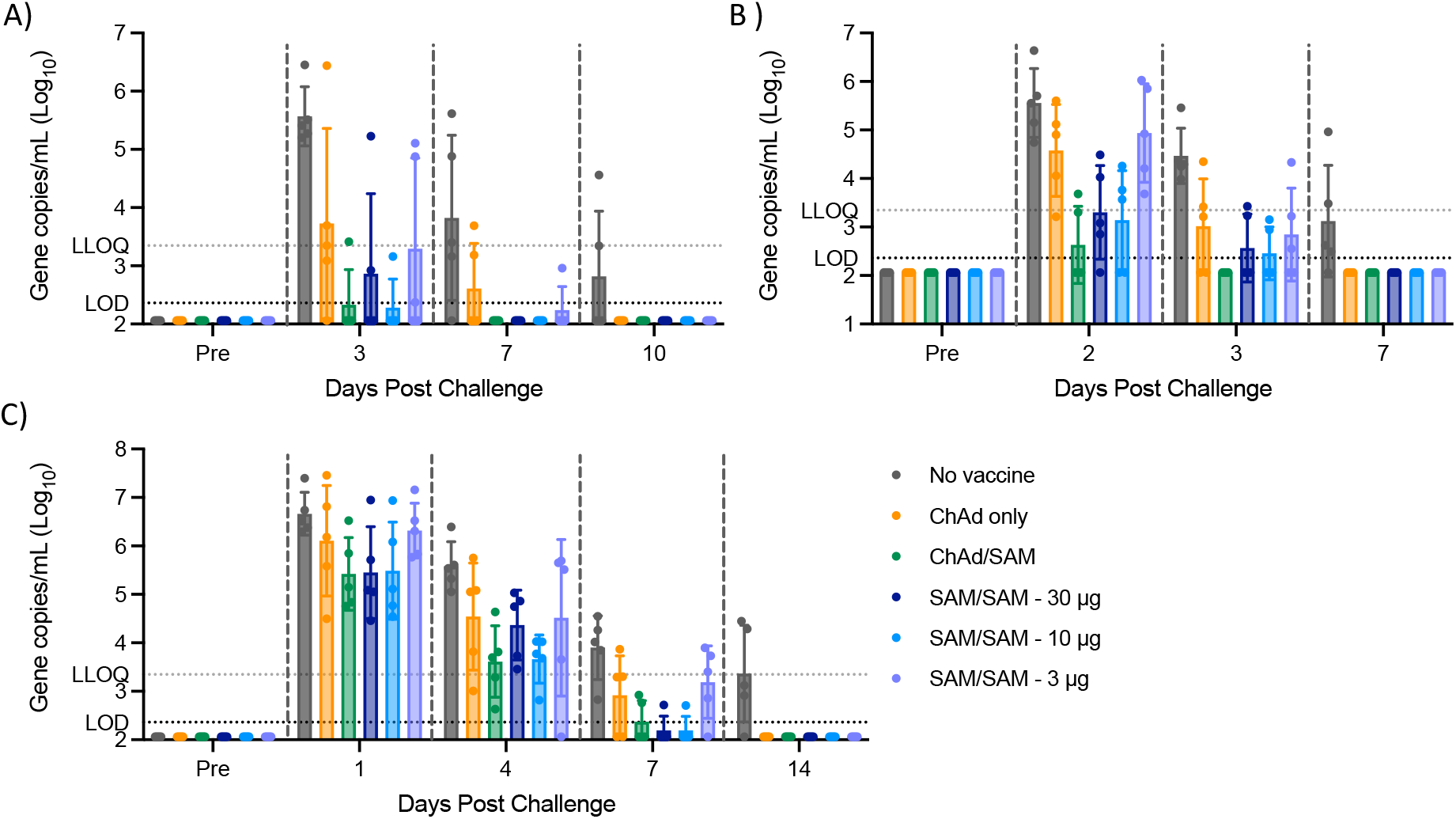
Total genomic RNA levels (N1) determined by RT-qPCR at specified timepoint post SARS-CoV-2 challenge for each animal in (A) bronchial alveolar lavage (B) oropharyngeal swab or (C) nasal swab. LOD = 230, samples below LOD set to ½ LOD. Geometric mean and SD.

**Supplemental Figure 7.**
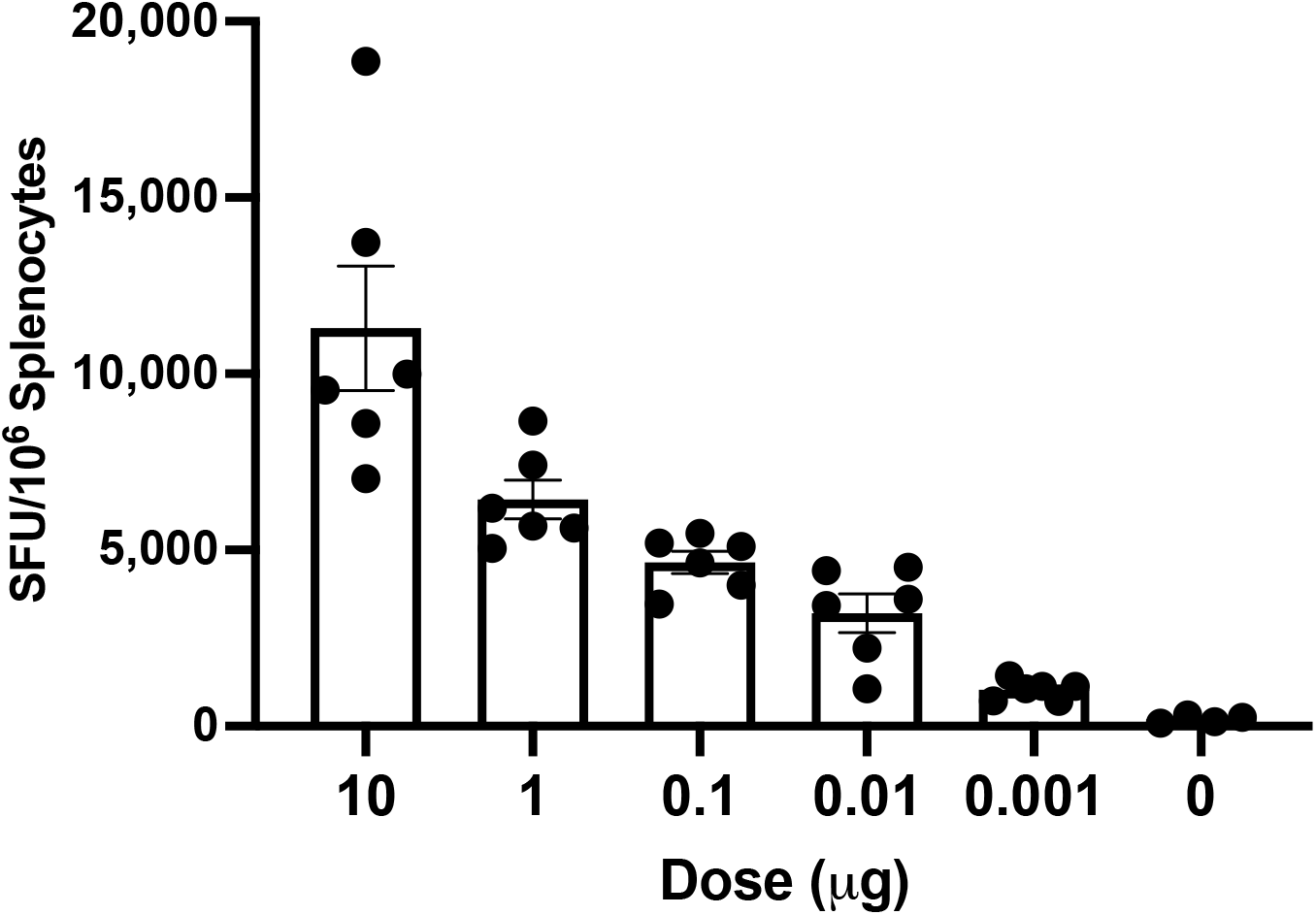
SAM induces potent T cell response at low doses in mice. Antigen-specific T cell response assessed by IFNγ ELISpot (sum of eight spike pools) in splenocytes in Balb/c mice (n=6/group) 2 weeks post immunization with SAM-Spike(V1) at the specified dose. Mean ± SE.

**Supplemental Figure 8.**
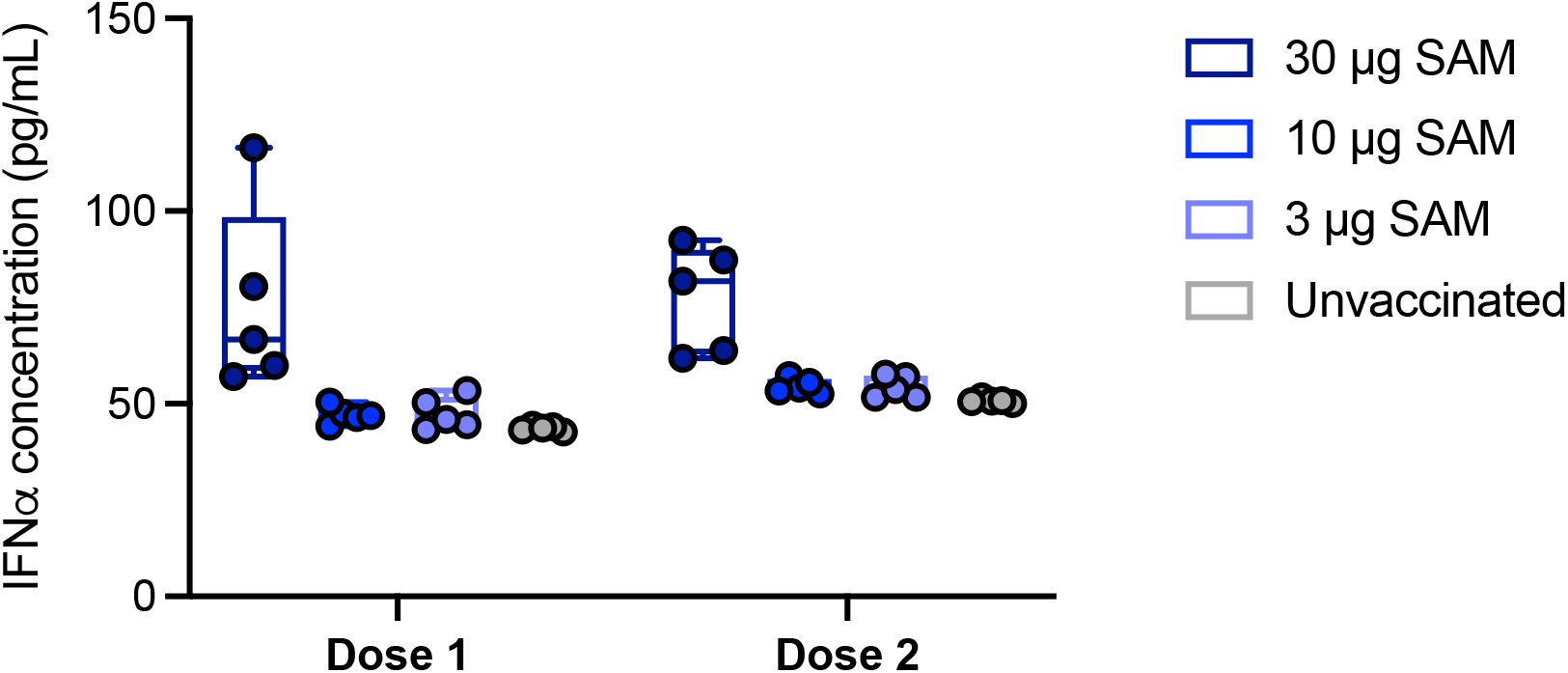
Serum IFN α2α concentration assessed by MSD in rhesus macaques (n=5/group) 8 hours post immunization with SAM-Spike(V2)-F2P at the specified dose. Median, IQR, range.

**Supplemental Figure 9.**
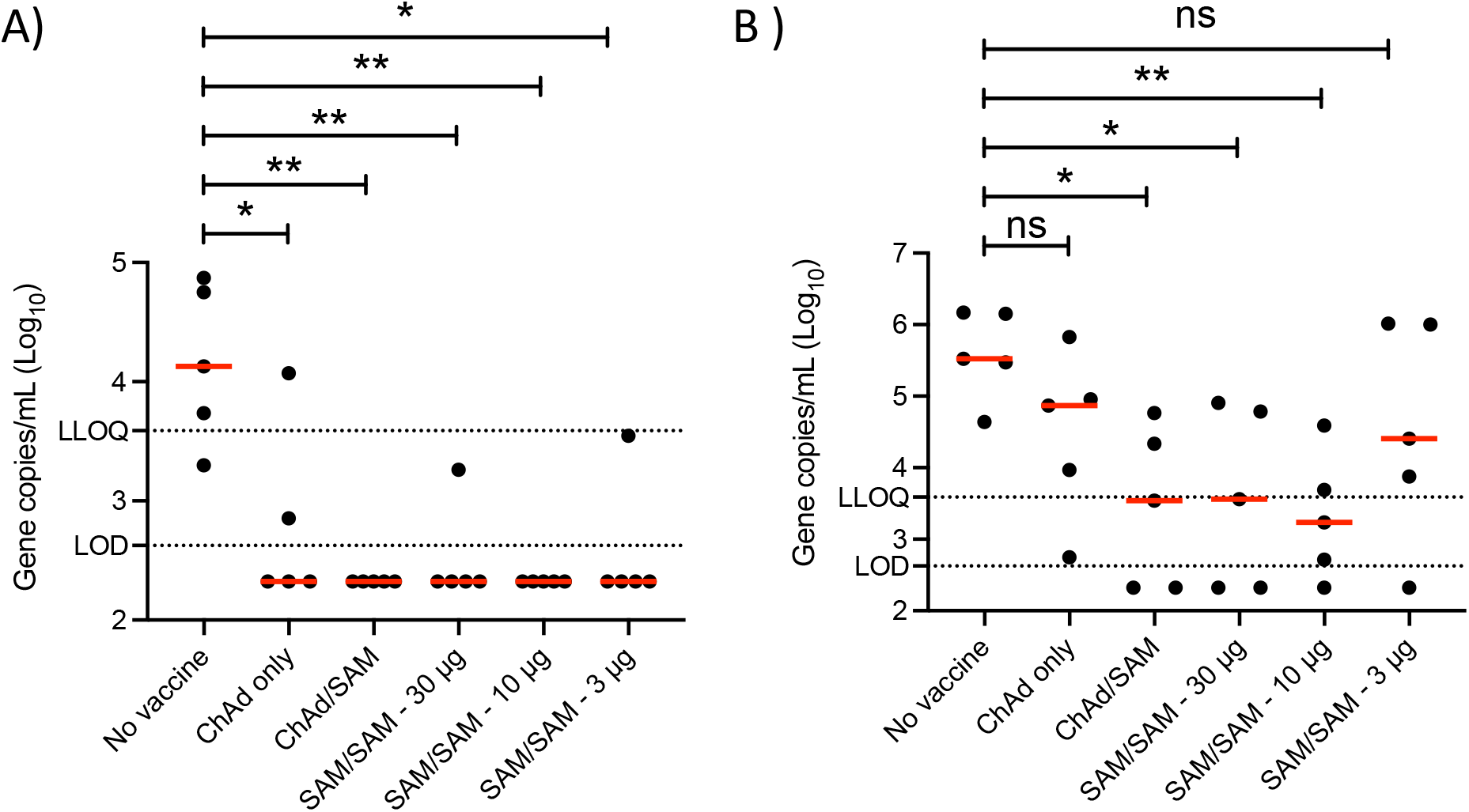
Viral replication is reduced in vaccinated NHP following SARS-CoV-2 challenge. (A) Peak subgenomic RNA levels in bronchial alveolar lavage for each animal determined by RT-qPCR between day 3 and day 10 post SARS-CoV-2 challenge. (B) Peak subgenomic RNA levels in nasal swab for each animal determined by RT-qPCR between day 2 and day 10 post SARS-CoV-2 challenge. LOD = 422, samples below LOD set to ½ LOD. LLOQ = 3,881. Red bar is median. Statistical analyses: Mann-Whitney test for each group compared to unvaccinated control group.

## References

1. Maxmen A. The fight to manufacture COVID vaccines in lower-income countries. Nature. 09 2021;597(7877):455–457. doi:10.1038/d41586-021-02383-z

2. El Sahly HM, Baden LR, Essink B, et al. Efficacy of the mRNA-1273 SARS-CoV-2 Vaccine at Completion of Blinded Phase. N Engl J Med. Sep 22 2021;doi:10.1056/NEJMoa2113017

3. Thomas SJ, Moreira ED, Kitchin N, et al. Safety and Efficacy of the BNT162b2 mRNA Covid-19 Vaccine through 6 Months. N Engl J Med. Sep 15 2021;doi:10.1056/NEJMoa2110345

4. Falsey AR, Sobieszczyk ME, Hirsch I, et al. Phase 3 Safety and Efficacy of AZD1222 (ChAdOx1 nCoV-19) Covid-19 Vaccine. N Engl J Med. Sep 29 2021;doi:10.1056/NEJMoa2105290

5. Sadoff J, Gray G, Vandebosch A, et al. Safety and Efficacy of Single-Dose Ad26.COV2.S Vaccine against Covid-19. N Engl J Med. 06 10 2021;384(23):2187–2201. doi:10.1056/NEJMoa2101544

6. Self WH, Tenforde MW, Rhoads JP, et al. Comparative Effectiveness of Moderna, Pfizer-BioNTech, and Janssen (Johnson & Johnson) Vaccines in Preventing COVID-19 Hospitalizations Among Adults Without Immunocompromising Conditions - United States, March-August 2021. MMWR Morb Mortal Wkly Rep. Sep 24 2021;70(38):1337–1343. doi:10.15585/mmwr.mm7038e1

7. Greinacher A, Selleng K, Palankar R, et al. Insights in ChAdOx1 nCov-19 Vaccine-induced Immune Thrombotic Thrombocytopenia (VITT). Blood. Sep 29 2021;doi:10.1182/blood.2021013231

8. Vogel AB, Kanevsky I, Che Y, et al. BNT162b vaccines protect rhesus macaques from SARS-CoV-2. Nature. 04 2021;592(7853):283–289. doi:10.1038/s41586-021-03275-y

9. Corbett KS, Flynn B, Foulds KE, et al. Evaluation of the mRNA-1273 Vaccine against SARS-CoV-2 in Nonhuman Primates. N Engl J Med. 10 15 2020;383(16):1544–1555. doi:10.1056/NEJMoa2024671

10. Vogel AB, Lambert L, Kinnear E, et al. Self-Amplifying RNA Vaccines Give Equivalent Protection against Influenza to mRNA Vaccines but at Much Lower Doses. Mol Ther. 02 07 2018;26(2):446–455. doi:10.1016/j.ymthe.2017.11.017

11. Atmar RL, Lyke KE, Deming ME, et al. Heterologous SARS-CoV-2 Booster Vaccinations: Preliminary Report. medRxiv. 2021:2021.10.10.21264827. doi:10.1101/2021.10.10.21264827

12. Schmidt T, Klemis V, Schub D, et al. Immunogenicity and reactogenicity of heterologous ChAdOx1 nCoV-19/mRNA vaccination. Nat Med. 09 2021;27(9):1530–1535. doi:10.1038/s41591-021-01464-w

13. van Doremalen N, Lambe T, Spencer A, et al. ChAdOx1 nCoV-19 vaccine prevents SARS-CoV-2 pneumonia in rhesus macaques. Nature. Jul 2020;doi:10.1038/s41586-020-2608-y

14. Pallesen J, Wang N, Corbett KS, et al. Immunogenicity and structures of a rationally designed prefusion MERS-CoV spike antigen. Proc Natl Acad Sci U S A. 08 2017;114(35):E7348–E7357. doi:10.1073/pnas.1707304114

15. Hsieh CL, Goldsmith JA, Schaub JM, et al. Structure-based Design of Prefusion-stabilized SARS-CoV-2 Spikes. bioRxiv. May 2020;doi:10.1101/2020.05.30.125484

16. McMahan K, Yu J, Mercado NB, et al. Correlates of protection against SARS-CoV-2 in rhesus macaques. Nature. Dec 2020;doi:10.1038/s41586-020-03041-6

17. Mercado NB, Zahn R, Wegmann F, et al. Single-shot Ad26 vaccine protects against SARS-CoV-2 in rhesus macaques. Nature. 10 2020;586(7830):583–588. doi:10.1038/s41586-020-2607-z

18. Rauch S, Gooch K, Hall Y, et al. mRNA vaccine CVnCoV protects non-human primates from SARS-CoV-2 challenge infection. bioRxiv. 2020:2020.12.23.424138. doi:10.1101/2020.12.23.424138

19. Green CA, Sande CJ, Scarselli E, et al. Novel genetically-modified chimpanzee adenovirus and MVA-vectored respiratory syncytial virus vaccine safely boosts humoral and cellular immunity in healthy older adults. J Infect. 05 2019;78(5):382–392. doi:10.1016/j.jinf.2019.02.003

20. Donovan-Banfield I, Turnell AS, Hiscox JA, Leppard KN, Matthews DA. Deep splicing plasticity of the human adenovirus type 5 transcriptome drives virus evolution. Commun Biol. 03 13 2020;3(1):124. doi:10.1038/s42003-020-0849-9

21. Stephenson KE, Le Gars M, Sadoff J, et al. Immunogenicity of the Ad26.COV2.S Vaccine for COVID-19. JAMA. 04 20 2021;325(15):1535–1544. doi:10.1001/jama.2021.3645

22. Pollock K, Cheeseman H, Szubert A, et al. Safety and immunogenicity of a self-amplifying RNA vaccine against COVID-19: COVAC1, a phase I, dose-ranging trial. Available at SSRN: https://ssrncom/abstract=3859294 or http://dxdoiorg/102139/ssrn3859294. 2021;

23. Sahin U, Muik A, Vogler I, et al. BNT162b2 vaccine induces neutralizing antibodies and poly-specific T cells in humans. Nature. 07 2021;595(7868):572–577. doi:10.1038/s41586-021-03653-6

24. Chu L, McPhee R, Huang W, et al. A preliminary report of a randomized controlled phase 2 trial of the safety and immunogenicity of mRNA-1273 SARS-CoV-2 vaccine. Vaccine. 05 12 2021;39(20):2791–2799. doi:10.1016/j.vaccine.2021.02.007

25. Low JG, de Alwis R, Chen S, et al. A phase 1/2 randomized, double-blinded, placebo controlled ascending dose trial to assess the safety, tolerability and immunogenicity of ARCT-021 in healthy adults. medRxiv. 2021:2021.07.01.21259831. doi:10.1101/2021.07.01.21259831

26. Zimmer C. Final trial results confirm that CureVac’s mRNA vaccine is far less protective than others. The New York Times. June 16, 2021. https://www.nytimes.com/2021/06/16/health/covid-vaccine-curevac.html

27. Bertoletti A, Le Bert N, Qui M, Tan AT. SARS-CoV-2-specific T cells in infection and vaccination. Cell Mol Immunol. 10 2021;18(10):2307–2312. doi:10.1038/s41423-021-00743-3

28. Wyllie D, Jones HE, Mulchandani R, et al. SARS-CoV-2 responsive T cell numbers and anti-Spike IgG levels are both associated with protection from COVID-19: A prospective cohort study in keyworkers. medRxiv. 2021:2020.11.02.20222778. doi:10.1101/2020.11.02.20222778

29. Bange EM, Han NA, Wileyto P, et al. CD8+ T cells contribute to survival in patients with COVID-19 and hematologic cancer. Nat Med. 07 2021;27(7):1280–1289. doi:10.1038/s41591-021-01386-7

30. Geers D, Shamier MC, Bogers S, et al. SARS-CoV-2 variants of concern partially escape humoral but not T-cell responses in COVID-19 convalescent donors and vaccinees. Sci Immunol. 05 25 2021;6(59)doi:10.1126/sciimmunol.abj1750

31. Tarke A, Sidney J, Methot N, et al. Impact of SARS-CoV-2 variants on the total CD4+ and CD8+ T cell reactivity in infected and vaccinated individuals. Cell Rep Med. Jul 20 2021;2(7):100355. doi:10.1016/j.xcrm.2021.100355

32. Bonifacius A, Tischer-Zimmermann S, Dragon AC, et al. COVID-19 immune signatures reveal stable antiviral T cell function despite declining humoral responses. Immunity. 02 09 2021;54(2):340-354.e6. doi:10.1016/j.immuni.2021.01.008

33. Catenacci DV, Liao C, Maron S, et al. Clinical outcomes and immune responses in a phase I/II study of personalized, neoantigen-directed immunotherapy in patients with advanced MSS-CRC, GEA and NSCLC. ESMO Congress 2021; 2021.

34. Chin JX, Chung BK, Lee DY. Codon Optimization OnLine (COOL): a web-based multi-objective optimization platform for synthetic gene design. Bioinformatics. Aug 01 2014;30(15):2210–2. doi:10.1093/bioinformatics/btu192

35. Scallan CD, Lindbloom JD, Tucker SN. Oral Modeling of an Adenovirus-Based Quadrivalent Influenza Vaccine in Ferrets and Mice. Infect Dis Ther. Jun 2016;5(2):165–83. doi:10.1007/s40121-016-0108-z

36. Geisbert TW, Lee AC, Robbins M, et al. Postexposure protection of non-human primates against a lethal Ebola virus challenge with RNA interference: a proof-of-concept study. Lancet. May 29 2010;375(9729):1896–905. doi:10.1016/S0140-6736(10)60357-1

37. Moodie Z, Price L, Gouttefangeas C, et al. Response definition criteria for ELISPOT assays revisited. Cancer Immunol Immunother. Oct 2010;59(10):1489–501. doi:10.1007/s00262-010-0875-4

38. Janetzki S, Price L, Schroeder H, Britten CM, Welters MJ, Hoos A. Guidelines for the automated evaluation of Elispot assays. Nat Protoc. Jul 2015;10(7):1098–115. doi:10.1038/nprot.2015.068

39. Hamilton MA, Russo RC, Thurston RV. Trimmed Spearman-Karber method for estimating median lethal concentrations in toxicity bioassays. Environmental Science & Technology. 1977/07/01 1977;11(7):714–719. doi:10.1021/es60130a004

40. Wölfel R, Corman VM, Guggemos W, et al. Virological assessment of hospitalized patients with COVID-2019. Nature. 05 2020;581(7809):465–469. doi:10.1038/s41586-020-2196-x

41. Corman VM, Landt O, Kaiser M, et al. Detection of 2019 novel coronavirus (2019-nCoV) by real-time RT-PCR. Euro Surveill. 01 2020;25(3)doi:10.2807/1560-7917.ES.2020.25.3.2000045

